# A crosstalk between E2F1 and GLP-1 signaling pathways modulates insulin secretion

**DOI:** 10.1101/2021.04.16.440172

**Authors:** Cyril Bourouh, Emilie Courty, Gianni Pasquetti, Xavier Gromada, Nabil Rabhi, Charlène Carney, Maeva Moreno, Raphaël Boutry, Laure Rolland, Emilie Caron, Zohra Benfodda, Patrick Meffre, Julie Kerr-Conte, François Pattou, Philippe Froguel, Amélie Bonnefond, Frédérik Oger, Jean-Sébastien Annicotte

## Abstract

Compromised β-cell function contributes to type 2 diabetes (T2D) development. The glucagon like peptide 1 (Glp-1) has emerged as a hormone with broad pharmacological potential toward T2D treatment, notably by improving β-cell functions. Recent data have shown that the transcription factor E2f1, besides its role as a cell cycle regulator, is involved in glucose homeostasis by modulating β-cell mass, function and identity. Here, we demonstrate a crosstalk between the E2F1, phosphorylation of retinoblastoma protein (pRb) and Glp-1 signaling pathways. We found that β-cell specific *E2f1* deficient mice (*E2f1*^β−/−^) presented with impaired glucose homeostasis and decreased glucose stimulated-insulin secretion mediated by exendin 4 (*i*.*e*., GLP1R agonist), which were associated with decreased expression of *Glp1r* encoding Glp-1 receptor (GLP1R) in *E2f1*^β−/−^ pancreatic islets. Decreasing E2F1 transcriptional activity with an E2F inhibitor in islets from nondiabetic humans decreased *GLP1R* levels and blunted the incretin effect of exendin 4 on insulin secretion. Conversely, overexpressing *E2f1* in pancreatic β cells increased *Glp1r* expression associated with enhanced insulin secretion mediated by GLP1R agonist. Interestingly, kinome analysis of mouse islets demonstrated that an acute treatment with exendin 4 increased pRb phosphorylation and subsequent E2f1 transcriptional activity. This study suggests a molecular crosstalk between the E2F1/pRb and GLP1R signaling pathways that modulates insulin secretion and glucose homeostasis.

## Introduction

Glucose homeostasis is finely tuned by the coordinated action of glucagon and insulin that are produced by pancreatic α and β cells, respectively. Dysregulation of this homeostatic process leads to type 2 diabetes (T2D), which is characterized by impaired function and mass of pancreatic β cells (1). Since lifestyle modifications are usually ineffective to cure T2D, pharmacological treatments are necessary to restore normoglycemia. However, most of the current therapeutic strategies are ineffective in controlling long-term glycemia, reinforcing the urgent need for new treatments.

Targeting the Glucagon Like Peptide-1 (Glp-1) pathway has demonstrated promising efficacy to treat T2D (2). Glp-1 is an enterohormone produced by intestinal L cells (3) and pancreatic α cells (4) that binds the Glp-1 receptor (GLP1R), a G protein-coupled receptor (GPCR) expressed by several cell types including the pancreatic β cells (5). The activation of this GPCR by natural Glp-1 or synthetic Glp-1 analogs, such as exendin-4 or Liraglutide (6) increases intracellular Ca^2+^ and activates adenylate cyclase to increase cyclic adenosine monophosphate (cAMP) levels and subsequent signaling pathways such as Protein kinase A (PKA) or Phosphoinositide 3-kinase (PI3K) (7). These mechanisms favor glucose-stimulated insulin secretion (GSIS) by pancreatic β cells (8, 9), inhibit glucagon secretion by pancreatic α cell (10) but also increase β-cell proliferation (11–13) and prevent pancreatic β-cell from apoptosis (14–16). Thus, GLP1R activation contributes to increased β-cell function and β-cell mass. We and others previously showed that E2F1-pRb-CDK4 pathway plays a major role to control β-cell mass and function (17–19). The transcription factor E2F1, the first member of the 8 E2F family genes (*i*.*e. E2F1* to *E2F8* (20)), binds the chromatin as an heterodimer with the Dimerization Partner-1 (DP-1) protein. The E2F1 transcriptional activity is finely regulated through its association with members of the retinoblastoma protein family such as pRb (21). During the G0/G1 phase of the cell cycle, the pRb/E2F1 complex at the chromatin inhibits E2F1 activity and the subsequent expression of E2F1 target genes (22, 23). When cells are stimulated by mitogenic signals such as growth factors, cyclin and cyclin-dependant kinase (CDK) complexes, such as CDK4/CyclinD1, phosphorylate pRb (24), leading to pRb release from E2F1, promoting E2F1 transcritptional activity and cell cycle progression. This mechanism is repressed through the activation of CDK inhibitors, namely p16^INK4A^, that bind to CDK and avoid cyclinD/CDK4 complex formation, and subsequently repress pRb phosphorylation and E2F1 transcriptional activity (25). Although these mechanisms mainly control the progression of the cell cycle (26), recent studies have reported important roles for the E2F1-pRb-CDK4 pathway beyond the *sole* regulation of cell proliferation (27, 28). Indeed, several reports have demonstrated the role of the E2F1 pathway in the control of metabolic functions in non-proliferative cells, including adipocytes (29, 30), hepatocytes (31–33), muscle and brown adipose tissue (34, 35) and pancreatic β cells (17–19, 36).

While E2F1 and Glp-1 pathways are both involved in the control of β-cell function and mass, whether they were physiologically connected remained unknown. Here, we demonstrate a crosstalk between the GLP1R and E2F1/pRb pathways in controlling insulin secretion. Using genetic mouse models in which the β-cell expression of *E2f1* is knockdown or the β-cell expression of *E2F1* is enhanced, we demonstrate that E2F1 modulates the Glp-1 signalling pathway through the transcriptional control of *Glp1r* expression by pancreatic β cells. Importantly, we found that a treatment of human islets with the pan E2F inhibitor HLM006474 decreases *E2F1* expression, as observed in type 2 diabetic human islets (37) and promoted a decrease in exendin-4 induced insulin secretion as well as *GLP1R* expression. Our results highlight a new molecular link between Glp-1 signaling and E2F1 in the control of β-cell function.

## Material & Methods

### Chemicals, antibodies and oligonucleotides

Chemicals, unless stated otherwise, were purchased from Sigma-Aldrich. Anti-phospho-Rb^S807/811^ (CS#9308) and anti-Rb 4H1 (CS#9309) antibodies were from Cell signaling. Anti-tubulin (T5168), anti-glucagon (G2654) and anti-HA antibodies were from Sigma-Aldrich, anti-insulin (A0564) was from DAKO. The oligonucleotides sequences used for various experiments are listed in Supplementary Table 1. The different plasmids used in this study were previously described (17). For ChIP-qPCR experiments, pCMV10 or pCMV10-hE2F1-Flag were used. The DNA sequence of the mouse *Glp1r* promoter corresponding to 1000 base pairs before the transcription start site was cloned in front of the luciferase gene (e-Zyvec).

**Table 1:** List of differentially phosphorylated peptides in mouse islets treated with 20 mM of glucose or 20 mM of glucose + 50 nM exendin-4.

| **Peptide_Name** | **Log2FC** | **-Log10(pValue)** |
| --- | --- | --- |
| CFTR_761_773 | 0,278437942 | 5,331759617 |
| F263_454_466 | 0,322237641 | 3,905287939 |
| KPB1_1011_1023 | 0,544231594 | 3,743610774 |
| KAP3_107_119 | 0,297545105 | 3,281205141 |
| CAC1C_1974_1986 | 0,439983994 | 3,19862641 |
| NFKB1_330_342 | 0,44198814 | 3,036551092 |
| ART_025_CXGLRRWSLGGLRRWSL | 0,491295815 | 2,829761112 |
| TY3H_65_77 | 0,45895353 | 2,78406994 |
| PTK6_436_448 | 0,476699829 | 2,766736962 |
| MYPC3_268_280 | 0,41120401 | 2,738905784 |
| CFTR_730_742 | 0,551521599 | 2,666052614 |
| EPB42_241_253 | 0,547589958 | 2,660160209 |
| STK6_283_295 | 0,241895363 | 2,599866013 |
| CREB1_126_138 | 0,454896599 | 2,505216277 |
| DESP_2842_2854 | 0,721435249 | 2,382496627 |
| TOP2A_1463_1475 | 0,27369532 | 2,337214199 |
| VTNC_390_402 | 0,364001602 | 2,311906038 |
| ADRB2_338_350 | 0,425349236 | 2,134341853 |
| RS6_228_240 | 0,351389259 | 2,120138342 |
| GRIK2_708_720 | 0,302527428 | 2,043075465 |
| ACM4_456_468 | 0,245583534 | 2,006511129 |
| REL_260_272 | 0,658538043 | 2,00029606 |
| PTN12_32_44 | 0,433181435 | 1,900544452 |
| KCNA6_504_516 | 0,30946508 | 1,898120619 |
| NCF1_321_333 | 0,422292382 | 1,862330501 |
| LIPS_944_956 | 0,571039379 | 1,832412796 |
| RYR1_4317_4329 | 0,268713325 | 1,739918105 |
| RAP1B_172_184 | 0,35980019 | 1,728827646 |
| P53_308_323 | 0,489605993 | 1,715349209 |
| STMN2_90_102 | 0,533314526 | 1,695191174 |
| SCN7A_898_910 | 0,412901074 | 1,677944879 |
| GBRB2_427_439 | 0,385804802 | 1,664691778 |
| GPSM2_394_406 | 0,662006855 | 1,657088411 |
| VASP_150_162 | 0,517954826 | 1,597883979 |
| NCF1_296_308 | 0,400952667 | 1,531899956 |
| ACM1_444_456 | 0,233731583 | 1,506976939 |
| CSF1R_701_713 | 0,468463272 | 1,496999651 |
| NMDZ1_890_902 | 0,388525337 | 1,444377647 |
| K6PL_766_778 | 0,431940705 | 1,423819681 |
| RB_803_815 | 0,63667804 | 1,366908432 |
| KCNA2_442_454 | 0,482689857 | 1,311541664 |
| KCNA3_461_473 | 0,427431911 | 1,307606387 |
| KAP2_92_104 | 0,262349129 | 1,301506585 |
| RAF1_253_265 | 0,372395277 | 1,260661604 |
| ANDR_785_797 | 0,144718647 | 1,206507993 |
| KIF2C_105_118_S106G | 0,34814772 | 1,197508754 |
| PLM_76_88 | 0,437398762 | 1,163264448 |
| VASP_271_283 | 0,316580296 | 1,159928639 |
| IF4E_203_215 | 0,299196482 | 1,157372437 |
| ACM5_494_506 | 0,158276245 | 1,084326059 |
| E1A_ADE05_212_224 | 0,267807961 | 1,073834338 |
| CDN1A_139_151 | 0,237322807 | 1,070748828 |
| MPIP1_172_184 | 0,359512806 | 1,006353783 |
| ERBB2_679_691 | 0,100586094 | 0,997823528 |
| BAD_112_124 | 0,551744461 | 0,982427985 |
| RB_242_254 | 0,238704041 | 0,955614557 |
| NOS3_1171_1183 | 0,18843396 | 0,946799825 |
| PLEK_106_118 | 0,226697609 | 0,932844109 |
| GPR6_349_361 | 0,281752437 | 0,925135001 |
| FRAP_2443_2455 | 0,36285305 | 0,921667898 |
| KAPCG_192_206 | 0,11823225 | 0,896310729 |
| NEK2_172_184 | 0,188770935 | 0,889058035 |
| ADDB_706_718 | 0,224111244 | 0,868373076 |
| GSUB_61_73 | 0,252330214 | 0,839803456 |
| RB_774_786 | 0,198471382 | 0,8389089 |
| ANXA1_209_221 | 0,396960884 | 0,83124898 |
| ESR1_160_172 | 0,294319004 | 0,778226067 |
| MPIP3_208_220 | 0,701477706 | 0,756986033 |
| PP2AB_297_309 | 0,193948269 | 0,714276568 |
| MBP_222_234 | 0,158284187 | 0,684763956 |
| PDPK1_27_39 | 0,57989502 | 0,683964096 |
| CENPA_1_14 | 0,196633816 | 0,654688779 |
| H32_3_18 | 0,243358299 | 0,652046094 |
| BRCA1_1451_1463 | -0,198002562 | 0,650452494 |
| CGHB_109_121 | 0,154372051 | 0,632171315 |
| FOXO3_25_37 | 0,100435257 | 0,614689704 |
| NR4A1_344_356 | 0,233428642 | 0,612047911 |
| KCNA1_438_450 | 0,328167111 | 0,607014033 |
| MP2K1_287_299 | 0,173790619 | 0,591064986 |
| MARCS_160_172 | 0,0965 | 0,577141439 |
| H2B1B_27_40 | 0,196655273 | 0,557926684 |
| ADDB_696_708 | 0,394567102 | 0,553493626 |
| KPCB_19_31_A25S | 0,150632858 | 0,546278049 |
| RADI_559_569 | -1,169470906 | 0,536950757 |
| VASP_232_244 | 0,715976059 | 0,510343188 |
| BAD_69_81 | 0,199600533 | 0,490755987 |
| GYS2_1_13 | 0,0941 | 0,489561781 |
| KS6A1_374_386 | 0,0465 | 0,480358957 |
| KPCB_626_639 | -0,312117904 | 0,414669067 |
| AKT1_301_313 | -0,326954275 | 0,397249622 |
| BAD_93_105 | 0,0751 | 0,379139344 |
| P53_12_24 | 0,220404223 | 0,369771968 |
| VIGLN_289_301 | 0,551927924 | 0,339256617 |
| CDK7_163_175 | 0,264778048 | 0,335196032 |
| NEK3_158_170 | 0,141193792 | 0,32813934 |
| RBL2_655_667 | 0,111330509 | 0,326777369 |
| NTRK3_824_836 | 0,236568034 | 0,322665551 |
| PDE5A_95_107 | 0,293000579 | 0,317922679 |
| BCKD_45_57 | -0,113957249 | 0,278838386 |
| KCNB1_489_501 | 0,162852213 | 0,275776027 |
| ERF_519_531 | 0,16679959 | 0,261561543 |
| TAU_524_536 | -0,220371976 | 0,198246162 |
| KCC2G_278_289 | 0,105276741 | 0,184374174 |
| PRKDC_2618_2630 | 0,0568 | 0,168460642 |
| RB_350_362 | -0,122649431 | 0,164536527 |
| COF1_17_29 | 0,274445355 | 0,164126941 |
| pTY3H_64_78 | 0,0457 | 0,160896065 |
| DCX_49_61 | 0,0896 | 0,126626152 |
| ELK1_356_368 | -0,128286898 | 0,125236389 |
| CDC2_154_169 | 0,0528 | 0,113166516 |
| MARCS_152_164 | 0,0287 | 0,101275344 |
| CD27_212_224 | 0,0273 | 0,097819167 |
| ACM1_421_433 | 0,0241 | 0,097117629 |
| CA2D1_494_506 | 0,0206 | 0,075288462 |
| LMNB1_16_28 | 0,0542 | 0,055812501 |
| ACM5_498_510 | 0,0227 | 0,033930329 |
| FIBA_569_581 | 0,0299 | 0,032636569 |
| PPR1A_28_40 | 0,0102 | 0,027891211 |
| PYGL_8_20 | -0,05 | 0,021152878 |
| pVASP_150_164 | 0,00465 | 0,012733952 |
| C1R_201_213 | -0,0088 | 0,008277163 |

### Animal experiments

Mice were maintained according to European Union guidelines for the use of laboratory animals. *In vivo* experiments were performed in compliance with the French ethical guidelines for studies on experimental animals (animal house agreement no. 59-350294, Authorization for Animal Experimentation, project approval by our local ethical committee no. APAFIS#2915-201511300923025v4). All experiments were performed with male mice. Mice were housed under a 12-hr light/dark cycle and given a regular chow. For high-fat diet (HFD) studies, 6-week-old mice were placed on a HFD (60% of calories from fat; Research Diet, D12492i) for 16 to 20 weeks. Oral glucose (OGTT) and intraperitoneal glucose and insulin tolerance tests (IPGTT and IPITT respectively) were performed as previously described (38) on 16 hours fasted animals for OGTT and IPGTT and 5 hours fasted animals for ITT. Glycemia was measured using the AccuCheck Performa (Roche Diagnostics). Circulating insulin levels were measured using the insulin ELISA kit (Mercodia). *E2f1* floxed (*E2f1*^*L2/L2*^) mice and rat insulin 2 promoter (RIP)-Cre mice were previously described (31, 39) and then further intercrossed to generate pure mutant *RIPcre*^*Tg/0*^*/E2f1*^*L2/l2*^ mice. A PCR genotyping strategy was subsequently used to identify RIPcre^+/+^::*E2f1*^*flox/flox*^ (*E2f1*^*β+/+*^) and RIPcre^Tg/+^::*E2f1*^*flox/flox*^ (*E2f1*^*β−/−*^) mice. Mice harboring the Rosa-26-loxP-LacZ-loxP-h*E2F1* conditional expression cassette (RFC mice) were obtained from Ulrike Ziebold (MDC, Berlin, Germany; (40)). Heterozygous RFC/+ mice were crossed with RIPcre^Tg/+^ mice to obtain RIPcre^Tg/+^::*RFC*^*Tg/+*^ (*E2f1*^*βover*^) and control RIPcre^+/+^::*RFC*^*Tg/+*^ (*E2f1*^*βCtrl*^) mice.

### Pancreatic islets studies

For mouse islets studies, pancreas was digested by type V collagenase (C9263, 1.5mg/ml) for 10 minutes at 37 °C as previously described (17, 38). After pancreas digestion and separation of pancreatic islets in a polysucrose density gradient medium, islets were purified by handpicking under a macroscope and were cultured during 16 hours before experiments. For GSIS experiments, approximately 30 islets were exposed to 2.8 mM glucose, 20 mM glucose or 20 mM glucose and 50 nM exendin-4 (Sigma, E7144) in Krebs-Ringer buffer supplemented with HEPES (Sigma, 83264) and 0.5% fatty-acid free BSA (Sigma, A7030). Insulin released in the medium was measured using the mouse insulin ELISA kit (Mercodia). Data are expressed as a ratio of total insulin content. Human pancreatic tissue was harvested from human, non-diabetic, adult donors. Isolation and pancreatic islet culture were performed as previously described (41). Human islets were treated for 48 hours with the pan-E2F inhibitor HLM006474 at 10µM (7). Data were expressed as a ratio of total insulin content. For mRNA quantification, human islets were isolated as described above and snap-frozen for further processing.

### Cell Culture, transfections and siRNA knock-down

Min6 cells (AddexBio) were cultured in DMEM (Gibco) with 15% fetal bovine serum, 100 mg/ml penicillin-streptomycin and 55 mM β-mercaptoethanol (Sigma, M6250). Cells were transfected with non-targeting siRNA mouse negative controls (siCont) and si*E2f1* (#13555, SMARTpool, Dharmacon) using Dharmafect1 (GE Dharmacon) and experiments were performed 48 hours later. Transient transfection experiments were performed in Min6 cells using Lipofectamine 2000 (Life Technologies) following the manufacturer’s instructions. Luciferase assays were performed 48 hours post-transfection and normalized to ß-galactosidase activity as previously described (17). For pharmacological treatments, Min6 cells and pancreatic islets were treated with HLM006474 (42) at 10μM for 48h. Exendin-4 (Sigma) was used at 50 nM at different time as indicated.

### Protein extracts and western blot experiments

Western blot was performed as previously described (43). Min6 cells were washed twice with cold PBS 1X and lysed with cell lysis buffer (50 mM Tris-HCl pH 8, 137 mM NaCl, 10% glycerol, 1% NP-40) supplemented with Protease Inhibitor cocktail (Roche) and phosphatase inhibitors (Thermofischer). Immunoblotting experiments were performed using 40 μg of total proteins and loaded on Precast SDS gel (Biorad). After electromigration, proteins were transferred on nitrocellulose membrane during 1h at 110V that were further incubated in TBS-Tween 0.05% (TBS-T) supplemented with 5% of milk. Membranes were incubated 16 h at 4 °C with primary antibodies as indicated in blocking buffer supplemented with 3% of bovine serum albumin or milk. After washing, membranes were incubated 1 h with the secondary antibody conjugated with horseradish peroxidase. The revelation of luminescent bands was performed using Pierce ECL Western blotting substrate or SuperSignal West Dura Extended duration substrate (ThermoFischer) with Chemidoc Xrs+ (Biorad).

### *In silico* analysis of the mouse *Glp1r* promoter region

The DNA sequence of the promoter region of the *Glp1r* gene was obtained from Ensembl (https://www.ensembl.org/index.html). Motif search was performed using LASAGNA-Search tool (44).

### Chromatin immunoprecipitation

Min6 cells were transfected with the empty vector pCMV10 or pCMV10-hE2F1-Flag. 48 hours after transfection, DNA-protein complexes from Min6 cells were formaldehyde-crosslinked to DNA at a final concentration of 1% for 10 minutes. The reaction was stopped by adding glycine at a concentration of 0.125M during 5 minutes. After cell lysis and sonication with the Bioruptor Pico (Diagenode, ref B01060010) for 8 minutes, proteins were immunoprecipitated with either the non-specific Ig G or anti-Flag 2 antibodies. After washing, protein-DNA complexes were decrosslinked by heating the samples for 16 hours at 65°C. DNA was purified using Mini Elute PCR purification kit (Qiagen) and qPCR were performed using promoter-specific primers.

### RNA extraction, measurements and profiling

Total RNA was extracted from Min6 cells using trizol reagent (Life Technologies). For mouse and human islets, total RNA was extracted with the RNeasy Plus Microkit (Qiagen) following manufacturer’s instructions. mRNA expression levels were measured after reverse transcription by quantitative real-time PCR (qRT-PCR) with FastStart SYBR Green master mix (Roche) using a LC480 instrument (Roche). qRT-PCR were normalized to cyclophilin mRNA levels. The results are expressed as the relative mRNA level of a specific gene expression using the formula 2^−ΔCt^.

### Kinome profiling

Serine-threonine kinase (STK) microarrays were purchased from PamGene International BV. Each array contained 140 phosphorylable peptides as well as 4 control peptides. Sample incubation, detection, and analysis were performed in a PamStation 12 according to the manufacturer’s instructions. The experiments were performed on mouse and human pancreatic islets as previously described (43).

### Immunohistochemistry and immunofluorescence

Immunofluorescence and immunohistochemistry were performed exactly as described previously (45). Pancreatic tissues were fixed in 10% formalin, embedded in paraffin and sectioned at 5 µm. For immunofluorescence microscopy analyses, after antigen retrieval using citrate buffer, 5-µm formalin-fixed paraffin embedded (FFPE) pancreatic sections were incubated with the indicated antibodies. Immunofluorescence staining was revealed by using a fluorescein-isothiocyanate-conjugated anti-mouse (for Glucagon and HA), fluorescein-isothiocyanate-conjugated anti-rabbit (for pRb^S807/811^), anti-guinea pig (for Insulin) secondary antibodies. Nuclei were stained with Hoechst.

### Statistical Analysis

Data are presented as mean ± s.e.m. Statistical analyses were performed using a two-tailed unpaired Student’s t-test, one-way analysis of variance (ANOVA) followed by Dunnett’s *post hoc* test or two-way ANOVA with Tukey’s *post hoc* tests comparing all groups to each other, using GraphPad Prism 9.0 software. Differences were considered statistically significant at p < 0.05 (*p < 0.05, ** p < 0.01, *** p < 0.001 and **** p < 0.0001).

## Results

### Loss of β cell-specific E2f1 expression impairs glucose tolerance under chow diet and during metabolic stress

To evaluate the role of *E2f1* in the control of glucose homeostasis, we bred *E2f1* floxed mice with Rip-Cre mice to generate pancreatic β cell-specific *E2f1*-deficient mice, named hereafter *E2f1*^β−/−^. The β-cell specific recombination efficiency was indeed confirmed through qRT-PCR analysis in islets isolated from *E2f1*^β−/−^ mice compared to *E2f1*^β+/+^controls (Figure 1A). In addition, *E2f1*^β+/+^ and *E2f1*^β−/−^ mice displayed comparable body weight (Figure 1B) and fasting glycemia (Figure S1A). To evaluate glucose homeostasis in those mouse models, we performed IPGTT in *E2f1*^β+/+^ and *E2f1*^β−/−^ mice fed with chow diet. Figure 1C and 1D shows that glucose clearance was impaired in *E2f1*^*β−/−*^ mice, associated with defects in secreting insulin in response to glucose injection (Figure 1E). Similarly, *E2f1*^β−/−^ mice demonstrated impaired glucose tolerance when challenged with an oral bolus of glucose compared to *E2f1*^β+/+^ mice (Figure 1F and 1G) associated with an alteration of insulin secretion (Figure 1H) Strikingly, although insulin levels were dampened in *E2f1*^β−/−^ mice, glucose-induced plasma Glp-1 levels were not affected by the deletion of E2f1 in β cells (Figure 1I). Because insulin secretion may be challenged in the context of insulin resistance, we next evaluated whether such metabolic dyshomeostasis promoted by β-cell selective *E2f1* knock-down could be exacerbated upon a metabolic stress such as high fat diet (HFD). Accordingly, we challenged controls and *E2f1*^*β−/−*^ mice with a HFD (60% of fat) for 16 weeks. Interestingly, despite similar body weight (Figure 1J and Supplementary figure S1B), glucose tolerance during IPGTT (Figures 1K and 1L), and insulin sensitivity (Supplementary figure S1C), insulin secretion was impaired in *E2f1*^β−/−^ mice as compared to controls fed a HFD (Figure 1M). Again, HFD-fed *E2f1*^β−/−^ mice demonstrated impaired glucose tolerance when challenged with an oral bolus of glucose compared to *E2f1*^β+/+^ mice (Figures 1N and 1O). Furthermore, insulin secretion was dampened 30 minutes after oral glucose intake in *E2f1*-deficient mice (Figure 1P), whereas circulating Glp-1 levels before and 10 minutes after an oral glucose bolus were not significantly different between control and *E2f1*^β−/−^ mice neither when fed a HFD (Figures 1Q). Altogether, our data demonstrate that the loss of *E2f1* expression within β cells impairs glucose homeostasis and insulin secretion upon physiologic conditions and during metabolic stress, despite the lack of differences in both body weight and Glp-1 circulating levels between controls and *E2f1*-deficient mice.

**Figure 1.**
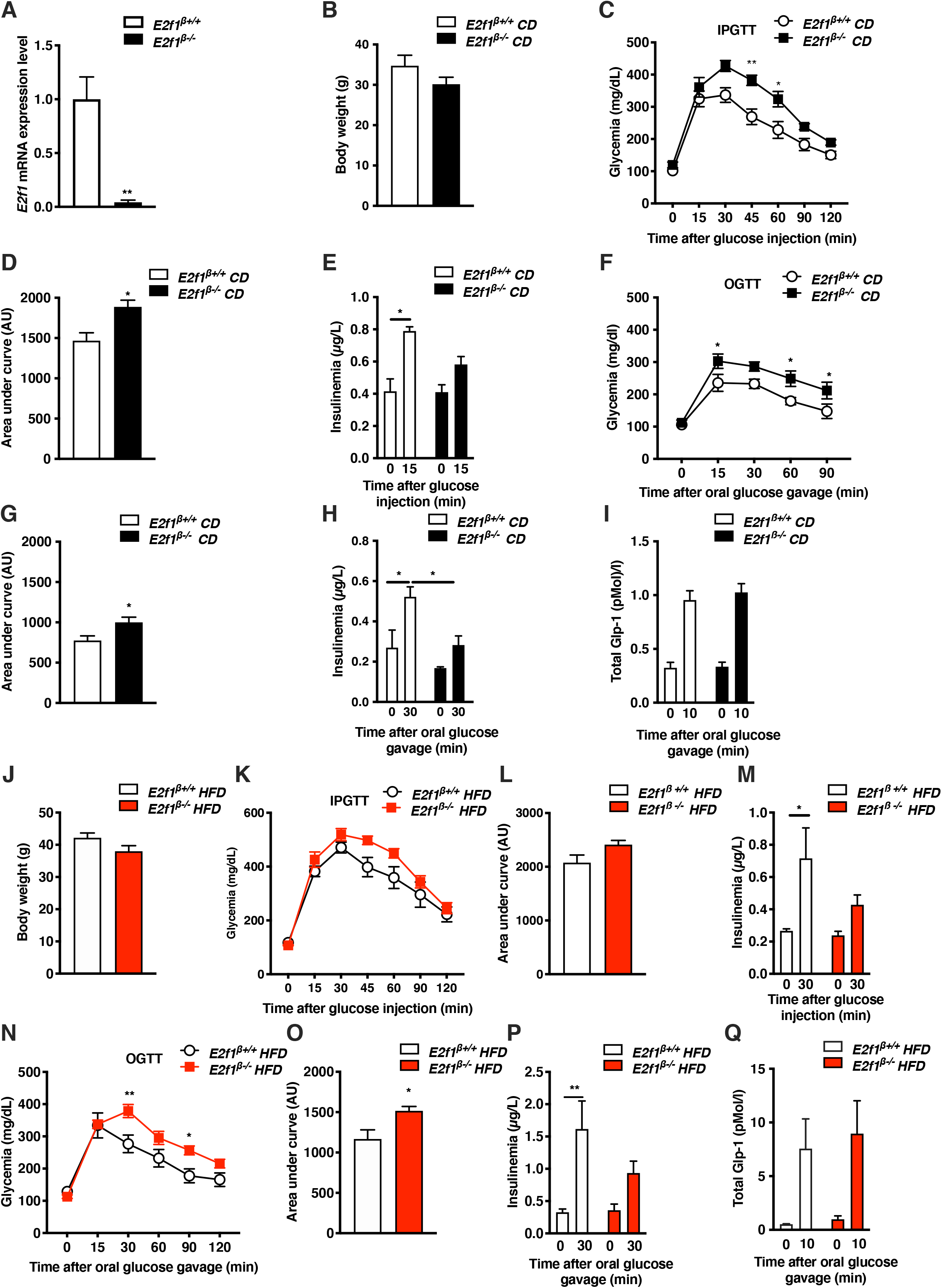
β cell-specific loss of *E2f1* expression impairs glucose tolerance under chow diet and during metabolic stress. **(A)** *E2f1* mRNA expression levels from pancreatic islets isolated from control (*E2f1*^β+/+^) and *E2f1*^β−/−^ mice (*n*=4). **(B)** Body weight of 16 week-old control (*E2f1*^β+/+^) and mutant (*E2f1*^β−/−^) mice fed with chow diet (CD, *n=4-6)*. **(C, D)** Intraperitoneal glucose tolerance test (IPGTT) in *E2f1*^β+/+^ and *E2f1*^β−/−^ mice fed with CD (*n*=5-7) and the corresponding area under curve (AUC, **D**). IPGTT was performed after 16h of fasting and glucose (2g/kg) was administrated by intraperitoneal injection. **(E)** Insulin plasma levels during IPGTT measured 0 and 15 minutes after glucose injection in CD fed *E2f1*^β+/+^ and *E2f1*^β−/−^ mice (*n*=5-7). **(F**,**G)** Oral glucose tolerance test (OGTT) in *E2f1*^β+/+^ and *E2f1*^β−/−^ mice fed with CD (*n*=4-6) and the corresponding area under curve (AUC, **G**). OGTT was performed after 16h of fasting and glucose (2g/kg) was administrated by oral gavage. **(H)** Insulin plasma levels during OGTT measured 0 and 30 minutes after glucose gavage in CD fed *E2f1*^β+/+^ and *E2f1*^β−/−^ mice (*n*=5-6). **(I)** Blood Glp-1 levels before and 10 minutes after an oral glucose gavage of *E2f1*^β+/+^ and *E2f1*^β−/−^ mice (n=5). **(J)** Body weight of *E2f1*^β−/−^ and *E2f1*^β+/+^mice fed a high fat diet (HFD) for 16 weeks (*n*=6). **(C**,**D)** IPGTT in *E2f1*^β+/+^ and *E2f1*^β−/−^ mice fed with HFD (*n*=6) and the corresponding area under curve (AUC, **D**). IPGTT was performed after 16h of fasting and glucose (1,5g/kg) was administrated by intraperitoneal injection. **(E)** Insulin plasma levels during IPGTT measured 0 and 30 minutes after glucose injection in HFD fed *E2f1*^β+/+^ and *E2f1*^β−/−^ mice (*n*=6). **(N**,**O)** OGTT **(N)** and its respective AUC (**O**, *n*=6). After 16h of fasting, OGTT (glucose 2g/kg) was performed on *E2f1*^β+/+^ and *E2f1*^β−/−^ mice fed a HFD for 16 weeks (*n*=6). **(P)** Insulin plasma levels during OGTT at 0 and 30 minutes following glucose challenge in *E2f1*^*β−/−*^ and *E2f1*^*β+/+*^ mice fed a HFD for 16 weeks (*n*=6). **(Q)** Total Glp-1 levels before and 10 minutes after an oral bolus of glucose in *E2f1*^*β+/+*^ and *E2f1*^*β−/−*^ mice after 20 weeks of HFD (*n*=7). All values are expressed as mean ± s.e.m. and were analyzed by two-tailed unpaired *t*-test (A, B, D, G, J, L, O) or two-way ANOVA followed by a Tukey’s *post-hoc* test (C, E, F, H, I, K, M, N, P, Q). *p < 0.05; **p<0.01.

### Overexpression of human E2F1 in murine β cells improve glucose homeostasis and insulin secretion upon metabolic stress

We next investigated whether *E2F1* overexpression in pancreatic β cells of wild type animals could mirror the phenotype observed in *E2f1*^*β−/−*^, *i*.*e*. increase insulin secretion in response to glucose. Therefore, we generated a mouse model that over-expresses the human form of *E2F1* (*hE2F1*) specifically in β cells (*E2f1*^***β****over*^). *Human E2F1* overexpression was confirmed at the mRNA levels (Figure 2A). The β-cell specific expression of hE2F1 proteins was confirmed using co-immunostaining with anti-insulin and anti-HA antibodies, an HA tag being fused with the *hE2F1* transgene (40) (Supplementary figure S2A). Overexpressing human E2F1 did not affect mouse *E2f1* transcript levels (Supplementary figure S2B). No particular phenotype was observed in these mice when fed with chow diet, including body weight (Figure 2B), fasting glycemia (Supplementary figure S2C), insulin sensitivity (Supplementary figure S2D), and intraperitoneal glucose tolerance (Figures 2C and 2D). Intriguingly, the insulin levels were increased in *E2f1*^***β****over*^ mice in response to *in vivo* glucose injection when compared to littermate controls (Figure 2E). Finally, *E2f1*^***β****over*^ mice demonstrated a significant improvement of glucose tolerance when challenged with an oral bolus of glucose (Figures 2F and G).

**Figure 2.**
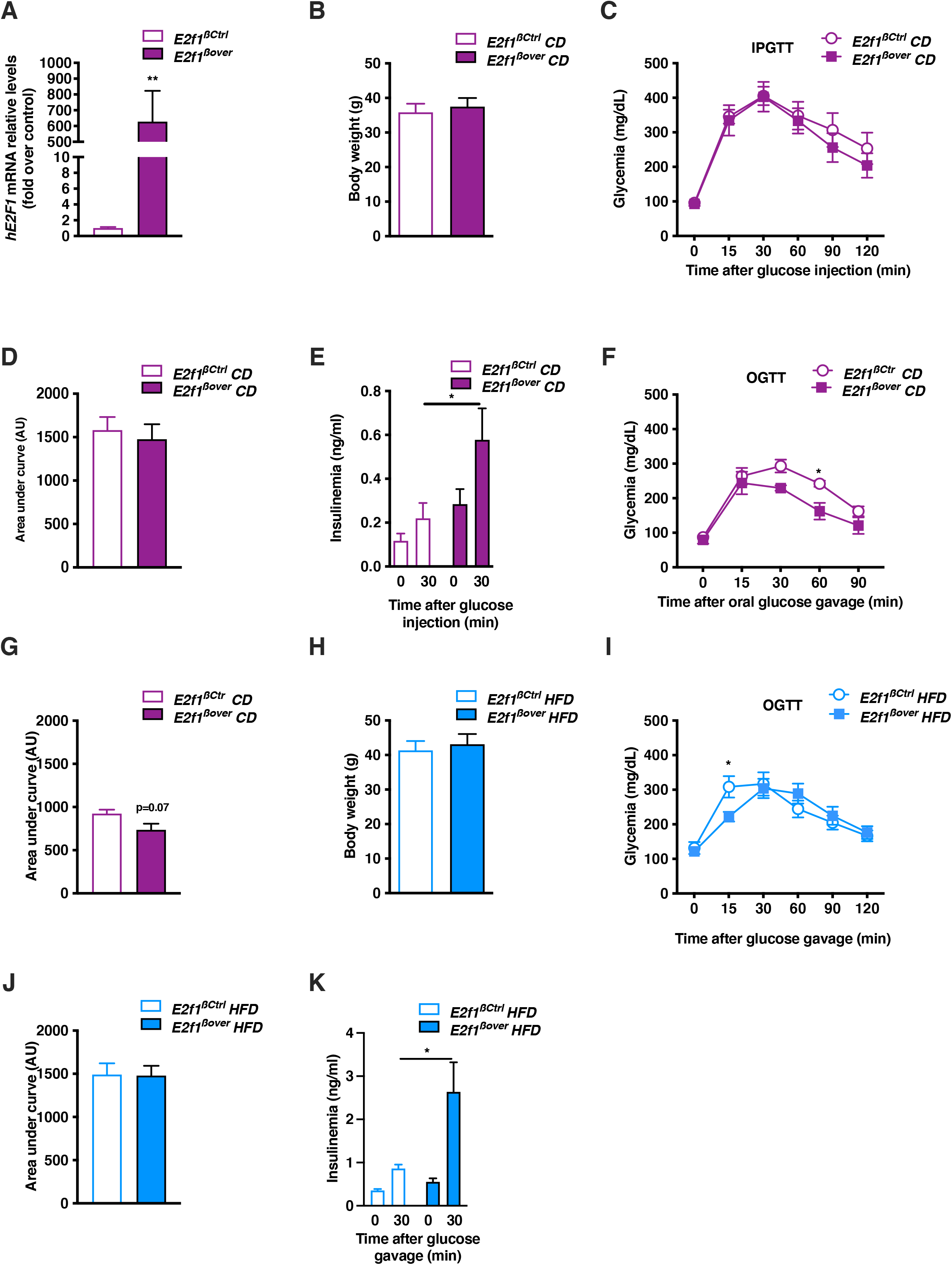
The overexpression of E2F1 in pancreatic β cells improves glucose homeostasis and insulin secretion under chow diet and during metabolic stress. **(A)** mRNA expression of human *E2F1* in pancreatic islets isolated from *E2f1*^βctrl^ and *E2f1*^βover^ mice (*n*=6). **(B)** Body weight of chow diet (CD) fed *E2f1* ^βctrl^ and *E2f1*^βover^ mice (*n*=10). **(C)** IPGTT was performed after 16h fasting and glucose was administrated by intraperitoneal injection (2g/kg) in *E2f1*^βctrl^ and *E2f1*^βover^ mice (*n*=11-15). **(D)** Area under curve (AUC) of IPGTT from *E2f1*^βctrl^ and *E2f1*^βover^ mice. **(E)** Insulin plasma levels during IPGTT before (0) and 30 minutes following glucose injection. (**F)** Blood glucose levels during OGTT of *E2f1*^βctrl^ and *E2f1*^βover^ mice under CD (*n*=5). (**G**) AUC of OGTT from *E2f1*^βctrl^ and *E2f1*^βover^ mice (*n*=5). **(H)** Body weight of *E2f1*^βctrl^ and *E2f1*^βover^ mice after 16 weeks of HFD (*n*=5). **(I)** OGTT was performed after 16h fasting and glucose was administrated by gavage (2g/kg) in *E2f1*^βctrl^ and *E2f1*^βover^ mice after 16 weeks of HFD (*n*=5). **(J)** AUC of OGTT from *E2f1*^βctrl^ and *E2f1*^βover^ mice (*n*=5). **(K)** Insulin plasma levels during OGTT at 0 and 30 minutes following glucose injection. All values are expressed as mean ± s.e.m. and were analyzed by two-tailed unpaired *t*-test (A, B, D, G, H, J) or two-way ANOVA followed by a Tukey’s *post-hoc* test (C, E, F, I, K). *p < 0.05; **p<0.01.

We next sought to evaluate the effect of h*E2F1* overexpression in pancreatic β cells on glucose homeostasis when challenged with HFD for 16 weeks. The body weight as well as the weight gain of *E2f1*^***β****over*^ mice remained comparable to control mice (Figure 2H and Supplementary figure S2E). When fed a HFD, overexpression of human *E2F1* in pancreatic β cells did not alter the insulin sensitivity (Supplementary Figure S2F) or the glucose tolerance after an intraperitoneal injection of glucose (Supplementary figures S2G and S2H). However, OGTT revealed an improved glucose tolerance in *E2f1*^***β****over*^ mice 15 minutes after glucose gavage (Figures 2I and 2J). Interestingly, *E2f1*^***β****over*^ mice secreted more insulin 30 minutes after an oral glucose load compared to control mice (Figure 2K). Altogether, our data demonstrate that, mirroring *E2f1*^β−/−^ mice, *E2f1*^***β****over*^ mice exhibit improved glucose homeostasis and insulin secretion upon physiologic conditions and during metabolic stress. Interestingly, our data highlight a differential outcome of β-cell specific *E2f1* modulation following IPGTT and OGTT raising the possibility that the incretin effects could be dependent on E2F1 levels within the β cell.

### Glucose-stimulated insulin secretion potentiation by the Glp-1 agonist exendin-4 is dependent on E2f1

To evaluate whether Glp-1 agonists modulate E2f1-dependent insulin secretion in a cell-autonomous manner, we performed static GSIS experiments in *E2f1* knockdown mouse Min6 β cell line and their controls. Strikingly, a 55% decrease in *E2f1* mRNA levels was sufficient to blunt GSIS (Figures 3A and 3B). Interestingly, while exendin-4 potentiated glucose effect on insulin secretion in control Min6 cells, that was not the case following *E2f1* knock-down (Figure 3B). To corroborate these findings, we performed a similar GSIS experiments using pancreatic islets isolated from *E2f1*^β+/+^ and *E2f1*^β−/−^ mice. Accordingly, whereas *E2f1*^*β+/+*^ pancreatic islets well responded to glucose, with an outcome potentiated in the presence of exendin-4, *E2f1*-deficient pancreatic islets failed to exhibit both enhanced response to glucose and potentiation by exendin-4 (Figure 3C). The treatment of non-diabetic human islets with the E2F pan inhibitor HLM006474 (42) provoked a marked decrease in insulin secretion in response to exendin-4 treatment, compared to control, DMSO-treated human islets (Figure 3D). Interestingly, GSIS assays on pancreatic islets isolated from control and *E2f1*^***β****over*^ mice showed that the overexpression of *E2F1* enhanced the potentiation impact of exendin-4 on glucose-induced insulin secretion (Figure 3E). Altogether, these data suggest that E2f1, both in mouse and human pancreatic islets, affects exendin-4 effects on GSIS.

**Figure 3.**
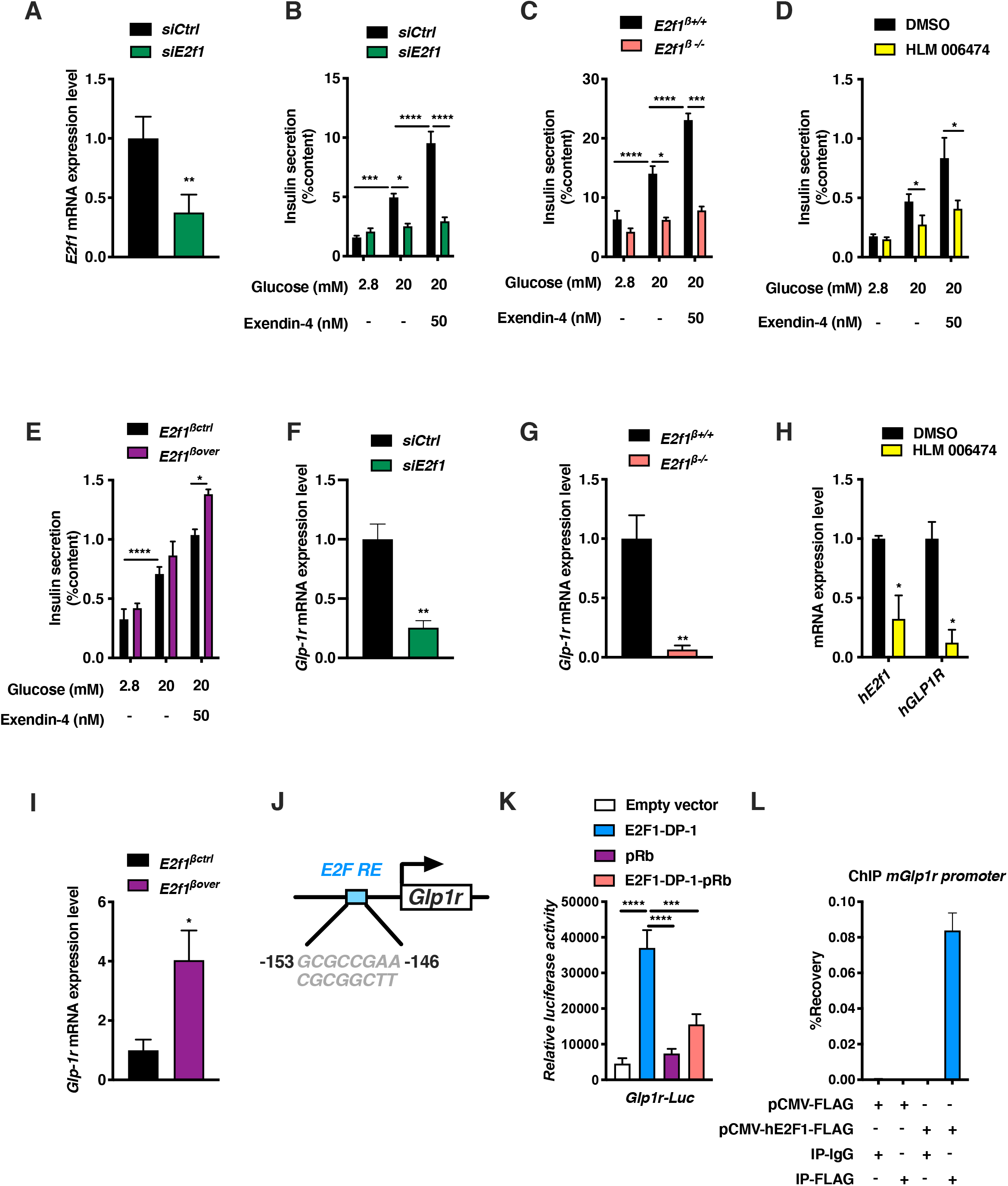
*E2f1* modulates Glp-1-mediated insulin secretion associated to *Glp1r* expression. **(A)** *E2f1* mRNA expression levels from control (*siCtrl*) or *E2f1* silencing (*siE2f1*) in Min6 cells (*n=5)*. **(B)** Glucose-stimulated insulin secretion (GSIS) measured in Min6 cells transfected with a *siCtrl* or *siE2f1* and exposed to 2.8 mM glucose, 20 mM Glucose or 20 mM Glucose + 50 nM exendin-4 (*n=7)*. **(C)** GSIS was measured in pancreatic islets isolated from *E2f1*^β+/+^ and *E2f1*^β−/−^ mice exposed to 2.8 mM glucose, 20 mM Glucose or 20 mM Glucose + 50 nM exendin-4 (*n=3-9)*. **(D)** GSIS experiment on human isolated islets treated for 48h with vehicle (0.1% DMSO) or 10µM HLM006474 and exposed to 2.8 mM glucose, 20 mM Glucose or 16.7 mM Glucose + 50 nM exendin-4 (*n=3)*. **(E)** GSIS measured in *E2f1*^βctrl^ and *E2f1*^βover^ isolated islets exposed to 2.8 mM glucose, 20 mM Glucose or 20 mM Glucose + 50 nM exendin-4 (*n=3-4)*. **(F)** *Glp1r* mRNA expression levels in *siCtrl* or *siE2f1* Min6 cells (*n=6)*. **(G)** *Glp1r* mRNA expression levels in *E2f1*^β+/+^ and *E2f1*^β−/−^ isolated islets (*n*=4). **(H)** *E2F1* and *GLP1R* mRNA expression levels in human islets treated for 48h with vehicle (DMSO 0,1%) or HLM006474 (10μM) (n=3). **(I)** *Glp1r* mRNA expression levels in *E2f1*^βctrl^ and *E2f1*^βover^ isolated islets (n=4-5). **(J)** Sequence of the E2F1 responsive element (E2F RE) identified in the mouse *Glp1r* promoter region. **(K)** Min6 cells were transiently co-transfected with an *mGlp1*r-promoter-luciferase construct (*Glp1r*-Luc vector) in the presence of a pcDNA3 vector (empty vector, negative control), E2F1-DP-1, pRb, E2F1-DP-1-pRb. Results were normalized to ß-galactosidase activity and are expressed as relative luciferase units. (K) ChIP-qPCR demonstrating E2F1 binding to the mouse *Glp1r* promoter from Min6 cells transfected with pCMV or pCMV-hE2F1-Flag plasmids. Chromatin from Min6 cells were incubated with IgG or Flag antibodies and qPCR were performed using primers amplifying the DNA region containing the E2F RE sequence found in the *Glp1r* promoter region (n=2). All values are expressed as mean ± s.e.m. and were analyzed by two-tailed unpaired *t*-test (A, F, G, H, I), one-way ANOVA followed by a Dunnett’s *post hoc* test (K) or two-way ANOVA followed by a Tukey’s *post-hoc* test (B, C, D, E). *p < 0.05; **p<0.01; ***p<0.001; ****p<0.0001.

### *E2f1 controls Glp1r* expression in pancreatic β cells

As E2f1 deficiency did not affect Glp-1 circulating levels (Figures 1I and 1Q) but inhibits insulin secretion in response to glucose stimulation and incretins, we then hypothesize that E2f1 modulates Glp-1 signaling in β cells rather than Glp-1 production. Since Glp-1 agonists modulate insulin secretion through the Glp1r (46) and our data support that β-cell response to exendin-4 depends on E2f1, we analyzed *Glp1r* expression in our different mouse and human models of *E2f1* modulation. Interestingly, the knockdown of *E2f1* in Min6 cells resulted in a significant decrease of *Glp1r* mRNA levels (around 65% of decrease, *p* = 0.0022; Figure 3F). We confirmed these results in *E2f1*^β−/−^ pancreatic islets (Figure 3G) and in human islets treated with the E2F inhibitor HLM006474 (Figure 3H). Interestingly, this E2F inhibitor not only decreased *GLP1R* expression in islets from nondiabetic humans, but also *E2F1* mRNA levels (Figure 3H). Notably, we observed a mirrored rise in *Glp1r* mRNA levels in isolated islets from *E2f1*^***β****over*^ mice (Figure 3I). *In silico* analysis revealed the presence of E2f1 binding sites in the promotor region of the *Glp1r* gene (Figure 3J and Supplementary figures S3A and S3B). To further demonstrate the ability of E2f1-dependent pathway to control *Glp1r* expression, we performed co-transfection experiments in Min6 cells. To monitor *Glp1r* transcriptional regulation, we generated a construct combining the mouse *Glp1r* promoter DNA sequence controlling the expression of the luciferase reporter gene. To activate or repress E2F1 pathways, we used expression vectors encoding E2F1 and its dimerization partner DP-1, as well as pRb, the *bona fide* repressor of E2F1(21). Our data demonstrate that activation of the E2F pathway by expressing the E2F1-DP1 heterodimer increased *Glp1r*-dependent luciferase activity while pRb repressed the increased luciferase activity mediated by E2F1-DP-1 (Figure 3K). Finally, we performed ChIP-qPCR experiments to determine whether E2F1 directly controls the *Glp1r* promoter at the chromatin level. In the absence of reliable antibodies to ChIP E2F1, Min6 cells were transfected with pCMV or pCMV-hE2F1-Flag to immunoprecipitate E2F1 using a Flag antibody. An enrichment at the *Glp1r* promoter was observed when cells were transfected with pCMV-hE2F1-Flag, demonstrating that E2F1 directly binds to the *Glp1r* promoter (Figure 3L). Altogether our data clearly highlight that E2F1 modulates *Glp1r* expression both in mouse and human islets, likely through the control of its promoter region. Strikingly, when E2F1-DP-1 are co-transfected with pRb, the activity of the *Glp1r* promoter is dampened (Figure 3K), showing that the activity of the E2F1-DP-1 complex on the Glp1r promoter is regulated by pRb.

### Glp-1 increases the phosphorylation of pRb through the activation of CDK signaling pathway in mouse and human pancreatic islets

Interestingly, the β cell-specific activation of the Glp-1 pathway by GLP1R agonist modulates the activity of several serine and threonine kinases and the subsequent phosphorylation of peptides controlling pancreatic β-cell functions (47). Since the molecular mechanisms of kinase activation controlled by the Glp-1 pathway are not fully understood, a kinome analysis using the Pamgene technology was performed on pancreatic islets isolated from *C57Bl6/J* mice treated with 20 mM glucose compared to 20 mM glucose plus 50 nM exendin-4 for 30 minutes (Figure 4A). Following these treatments, an increase in 43 phosphorylation sites of 41 peptides was observed when pancreatic islets where treated with glucose-exendin-4 compared to glucose alone (Figure 4B and table 1). It is noteworthy that treatment of mouse islets with 20 mM glucose plus 50 nM exendin-4 increased Creb phosphorylation at Serine 133 (Creb^S133^, Figure 4B, *Log2 Fold Change* = 0.45, *p* = 0.003), as previously shown (48). Strikingly, the co-treatment of mouse pancreatic islets with 20 mM glucose plus 50 nM exendin-4 for 30 minutes increased the phosphorylation level of the pRb on serine 807/811 (pRb^S807/811^; *Log2 Fold Change* = 0.63, *p* = 0.043), the *bona fide* repressor of E2F1 transcriptional activity. The exendin-4-mediated induction of pRb^S807/811^ phosphorylation was further confirmed in the mouse Min6 β cells through western blotting and immunofluorescence analysis (Supplementary figures S4A, S4B and S4C). Integrating the phosphorylation data using the Bionavigator software developed by Pamgene revealed the specific activation of kinases upon exendin-4 treatment, including several CDKs (Figure 4C). Finally, ingenuity pathway analysis (IPA) further demonstrated that several canonical pathways were controlled by exendin-4 treatment, such as the signaling pathways related to opioid, AMPK or PKA (Figure 4D). Altogether, these results demonstrate that the activation of the Glp-1 pathway by its agonist exendin-4 in mouse pancreatic islets activates several kinases, including CDKs, leading to the phosphorylation of pRb.

**Figure 4.**
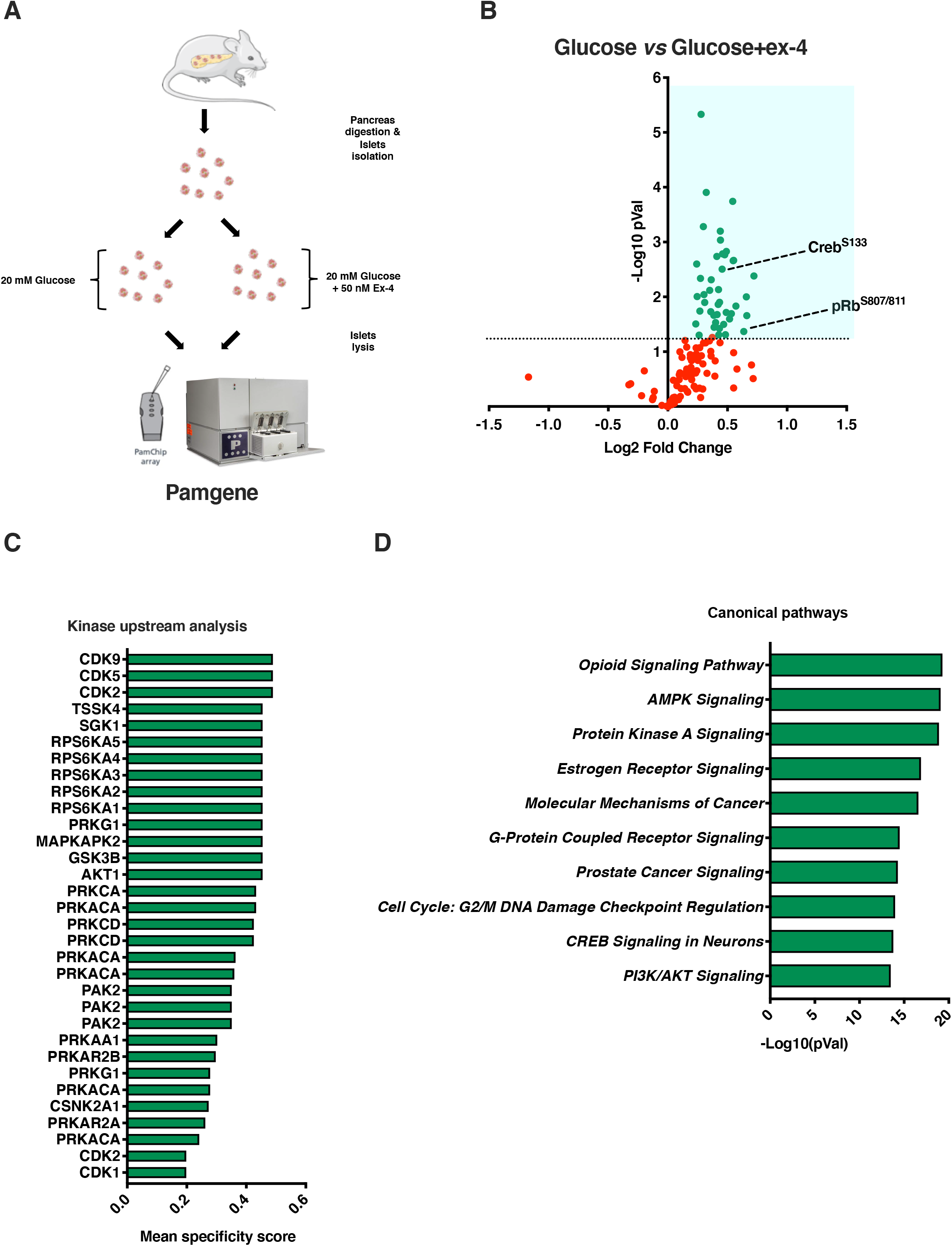
Short term Glp1r activation induces pRb phosphorylation in pancreatic islets. **(A)** Schematic representation of the kinome profiling strategy following exendin-4 treatment of mouse pancreatic islets. **(B)** Volcano plot exhibiting differential peptide phosphorylation in mouse pancreatic isolated islets treated with 20mM glucose (*n*=3) or 20 mM glucose + 50 nM exendin-4 (*n*=3) during 30 minutes. Results were obtained using the Pamgene technology and are expressed as Log2 Fold Change and –Log10 (pvalue) comparing differential phosphorylation levels between 20mM glucose and 20 mM glucose + 50 nM exendin-4. **(C)** Analysis of upstream kinases involved in differential peptide phosphorylation using the Bionavigator Software. **(D)** Differential peptide phosphorylation was analyzed using Ingenuity Pathway Analysis (IPA) and identified canonical pathways controlled by exendin-4 treatment. All values are expressed as mean ± s.e.m. and were analyzed by two-tailed unpaired *t*-test (B).

### Exendin-4 increases E2F1 transcriptional activity

These data led us to postulate that GLP1R activation through an exendin-4 treatment could increase E2F1 transcriptional activity, since the *E2f1* target gene expression is strongly controlled by the phosphorylation status of pRb (21). To test this hypothesis, we first performed luciferase assays using a plasmid containing E2F responsive elements (E2F-RE) cloned in front of the Thymidine Kinase (Tk) promoter and the luciferase gene (E2F-RE-Tk-Luc). Upon treatment with 2.8 mM glucose, the co-transfection with the E2F1 and DP-1 expression vectors induced luciferase activity compared to control conditions, validating increased E2F1 transcriptional activity in the presence of E2F1 and DP-1 (Figure 5A). The treatment of Min6 cells with 20 mM glucose further increased E2f1 transcriptional activity when compared to 2.8 mM glucose treatment (Figure 5A). Interestingly, this effect was even more potentiated with a 50 nM exendin-4 treatment (Figure 5A). Altogether, these results suggest that exendin-4 enhanced the transcriptional activity of E2f1 on the promoter of its target genes. On the other hand, the co-transfection with a pRb expression vector nearly abolished E2f1 transcriptional activity (Figure 5A), thus demonstrating as expected, that pRb inhibits E2f1 transcriptional activity. However, when Min6 cells were treated with 50 nM of exendin-4, the pRb repressive effect was partially alleviated, thus allowing a weak transcriptional activity of the E2F1-DP1 heterodimer on the E2F-RE-Tk promoter (Figure 5A). These data then suggest that a loss of the pRb repressive activity following the treatment with the GLP1R agonist (Figure 5A). Since exendin-4 increased E2f1 transcriptional activity, we next wondered whether the expression of genes involved in β-cell mass and/or function was modulated upon exendin-4 treatment. Since PDX-1, a master regulator of β-cell identity that controls *Insulin* gene expression (49) is regulated by exendin-4 treatment (50, 51), we checked whether short term treatment with 50nM of exendin-4 (1 to 4 hours) modulated its expression in Min6 cells or mouse pancreatic islets in which *E2f1* expression level is downregulated. Our results showed that treatment with 50 nM of exendin-4 increased the mRNA levels of *E2f1* (Figure 5B) and *Pdx1* (Figure 5C) in Min6 cells transfected with a non-targeting, control siRNA. Knocking down *E2f1* in Min6 cells completely blocked the exendin-4 mediated raise in *E2f1* (Figure 5B) and *Pdx-1* (Figure 5C) mRNA levels. These results were also observed when C57Bl6J pancreatic islets were treated with glucose and exendin-4 (Figures 5D and 5E). Interestingly, treating *E2f1*^β−/−^ pancreatic islets with exendin-4 failed to increase *E2f1*, confirming the reproducibility and robustness of our model, but most interestingly failed to increase *Pdx1* expression levels (Figures 5F and 5G, respectively). Altogether, these results suggest that exendin-4 modulates pRb phosphorylation and E2F1 transcriptional activity, which in turn modulates insulin secretion through the upregulation of E2f1 dependent genes such as *E2f1* itself or *Pdx1* mRNA levels.

**Figure 5.**
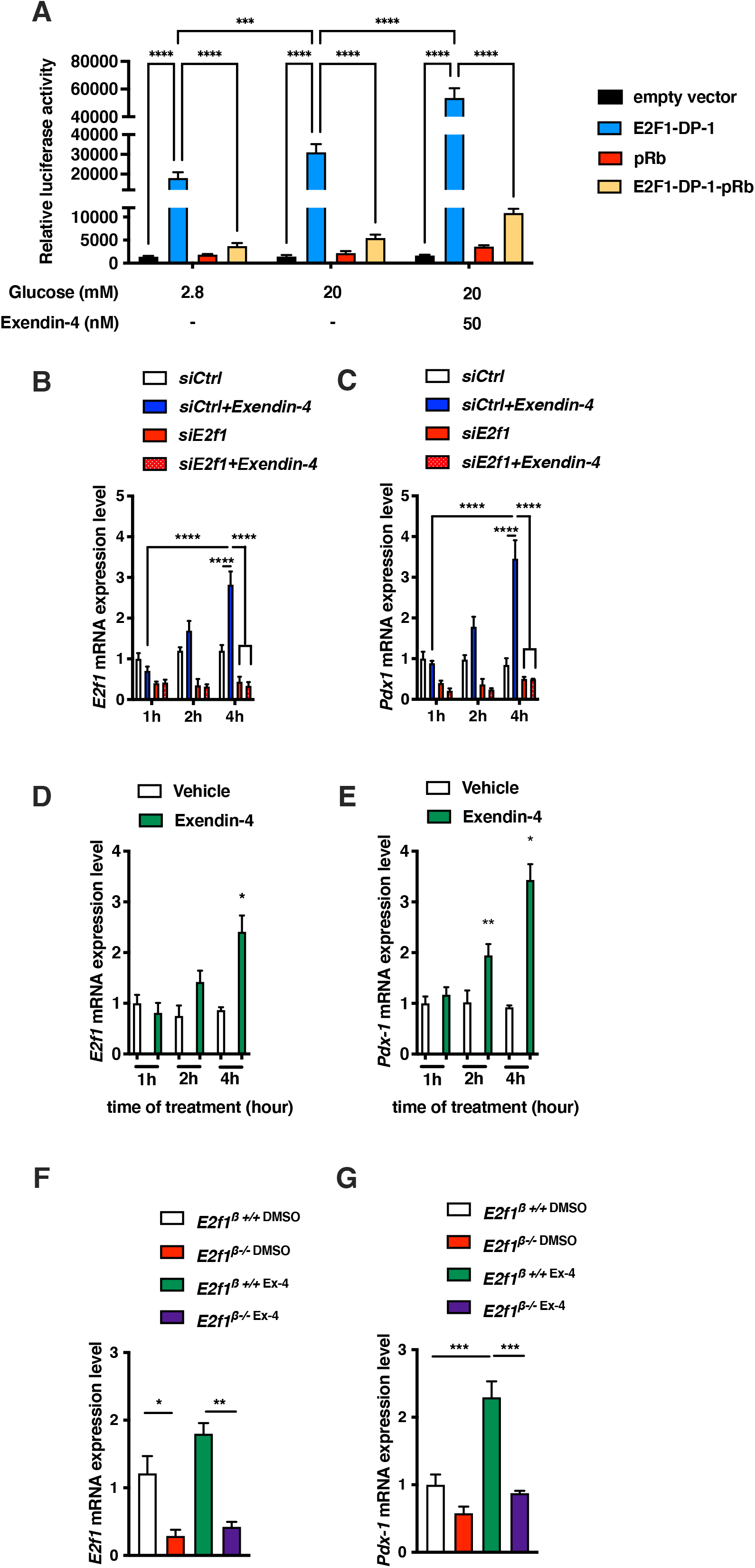
Exendin-4 treatment modulates E2f1 transcriptional activity. **(A)** Min6 cells were transiently co-transfected with an E2F-RE-Tk-luciferase construct (E2F-RE-Tk-Luc vector) in the presence of the empty pcDNA3 vector (negative control), E2F1-DP-1, pRb or E2F1-DP-1-pRb, and treated during 30 minutes with 2.8 mM glucose, 20mM glucose or 20 mM glucose + 50 nM exendin-4 (*n*=3). (**B**) *E2f1* mRNA expression from Min6 control (*siCtrl*) or E2f1 silenced (*siE2f1*) cells treated with 20mM glucose (Gluc) or 20mM glucose and 50nM exendin-4 (Gluc+Ex4) during 1, 2 or 4 hours (*n*=3). (**C**) Pdx1 mRNA expression from *siCtrl* or *siE2f1*-Min6 cells treated with 20mM glucose (Gluc) or 20mM glucose and 50nM exendin-4 (Gluc+Ex4) during 1, 2 or 4 hours (*n*=3). (**D**) *E2f1* mRNA expression from pancreatic islets isolated from 12-week-old C57Bl6J mice and treated with 20mM glucose (Gluc) or 20mM glucose and 50nM exendin-4 (Gluc+Ex4) during 1, 2 or 4 hours (*n*=3). **(E)** *Pdx1* mRNA expression from pancreatic islets isolated from 12-week-old C57Bl6J mice and treated with 20mM glucose (Gluc) or 20mM glucose and 50nM exendin-4 (Gluc+Ex4) during 1, 2 or 4 hours (*n*=3). **(F)** mRNA expression of *E2f1* in pancreatic islets isolated from *E2f1*^β−/−^ and *E2f1*^β+/+^ mice treated for 4 hours with 20mM glucose (Gluc) or 20mM glucose and 50nM exendin-4 (Gluc+Ex4) (*n*=3-4). **(G)** mRNA expression of *Pdx1* in pancreatic islets isolated from *E2f1*^β−/−^ and *E2f*^1 β+/+^mice treated for 4 hours with 20mM glucose (Gluc) or 20mM glucose and 50nM exendin-4 (Gluc+Ex4) (*n*=3-4). All values are expressed as mean ± s.e.m. and were analyzed by two-way ANOVA followed by a Tukey’s *post-hoc* test (A, B, C, D, E, F, G). *p < 0.05; **p<0.01; ***p<0.001.

## Discussion

Here, we report that E2F1 transcription factor, primarily known to control cell cycle progression (52), plays a key role in pancreatic β-cell function through the control of Glp-1 signaling, both in mice and human islets. Following previous studies demonstrating that E2F1-pRb pathway controls pancreatic β-cell mass (18, 36, 53, 54) and function (17, 55), we identify here a new role for this transcription factor linking this pathway to the druggable Glp-1 signaling. Indeed, we demonstrate that the specific loss of *E2f1* expression in β cells leads to impaired oral glucose tolerance, both under chow and upon metabolic stress, associated to a loss of insulin secretion in response to glucose, despite normal Glp-1 circulating levels. Interestingly, we have also shown that human *E2F1* overexpression in mouse pancreatic β cells counteracts glucose impairment upon metabolic stress by inducing insulin secretion associated with a raise in *Glp1r* mRNA levels. In this study, we also report a crosstalk between Glp-1 and E2F1-pRb signaling pathways, where the Glp1r agonist exendin-4 modulates pRb phosphorylation status, both in mice and human islets, and subsequent E2f1 transcriptional activity (Figure 6).

**Figure 6.**
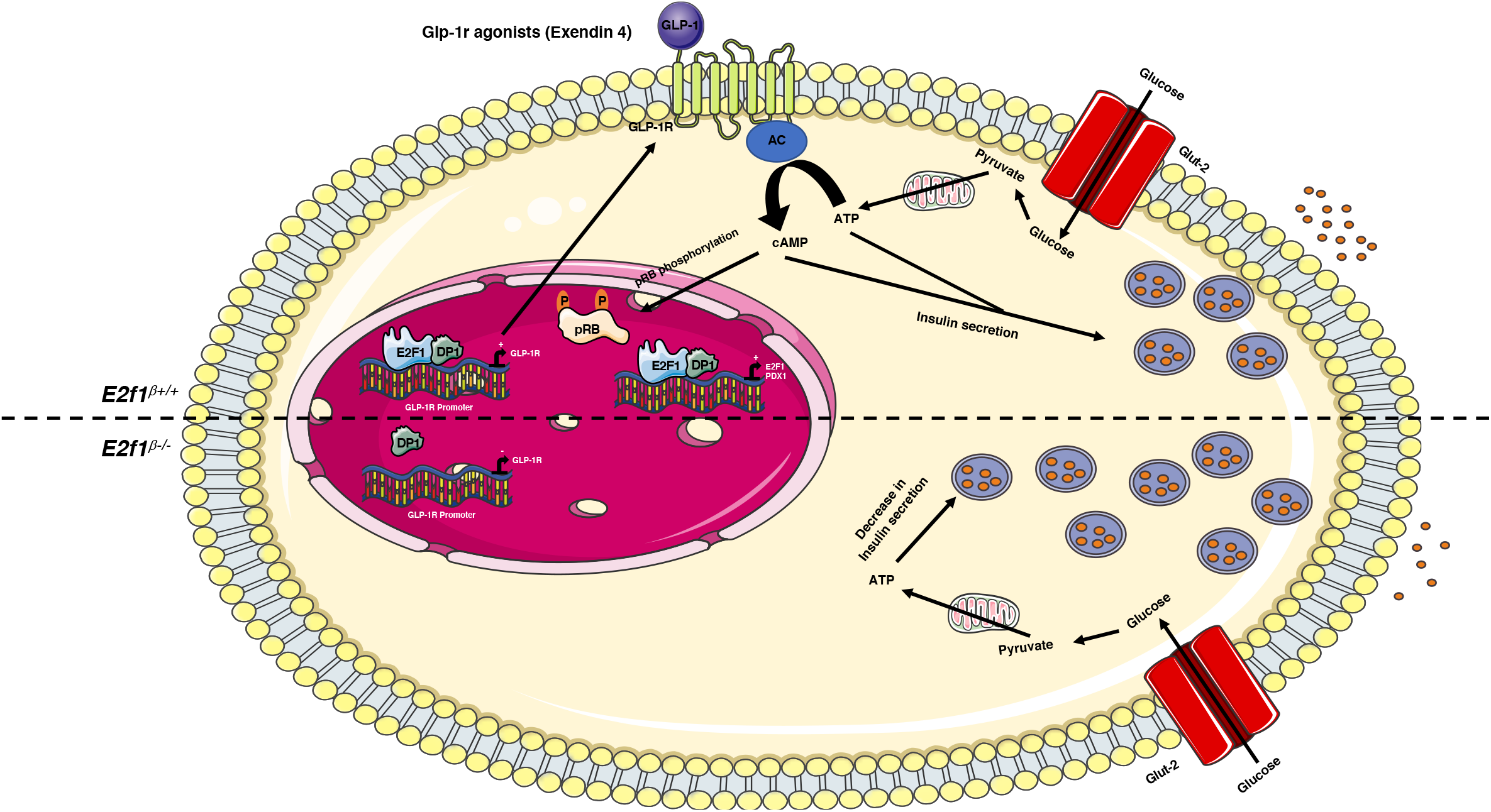
A model linking E2f1 and GLP-1 signaling pathways in pancreatic β cell. In pancreatic β cells, the transcription factor E2f1 binds to the *Glp1r* promoter and positively modulates its expression. In addition, glucose entry in cells through the Slc2a2 (Glut-2) transporter increases the level of ATP and stimulates insulin secretion. Glp1r activation by exendin-4 potentiates glucose-stimulated insulin secretion. In addition, exendin-4 increases pRB phosphorylation, E2f1 transcriptional activity as well as the expression of *E2f1* and *Pdx-1*. In *E2f1*^β−/−^ pancreatic islets, *Glp-1r* expression is decreased, resulting in a loss of potentiation effects of Exendin-4 on GSIS. AC: Adenylate cyclase; ATP: Adenosine tri-phosphate; cAMP: cyclic Adenosine mono-phosphate; GLP-1: Glucagon-like peptide-1; GLP-1R: Glucagon-like peptide-1 receptor; GLUT-2: Glucose transporter 2; E2F1: E2F transcription Factor 1; DP1: Dimerization Partner 1; pRB: Retinoblastoma protein.

Along the last 2 decades, a particular emphasis has been put toward the development of specific GLP1R agonists to treat T2D (56). Indeed, GLP1R agonists improve glycaemia in patients with T2D by increasing insulin synthesis, secretion and by inducing β-cell proliferation (7, 57). Although targeting the GLP1R signaling pathway is a promising therapeutic strategy, it has been reported however that the incretin effects decrease in humans with the onset of obesity and T2D, thus urging us to uncover the underlying mechanism involved in the loss of GLP1R agonist sensitivity in obese and T2D patients (58, 59). Several studies suggest that part of this deleterious mechanism could rely on the downregulation of the *Glp1r* expression in different tissues involved in metabolic homeostasis, including pancreatic β cells (60, 61). Indeed, Kimura et al demonstrated that obesity induces a loss of *Glp1r* expression in vascular tissue (62). Other studies have demonstrated that hyperglycemia not only downregulates both *Glp-1* et *Gip* receptors expression in pancreatic islets but also impairs Glp-1 signaling pathway through the loss of *Glp1r* at the β-cell cytoplasmic membrane (63). Then, preserving *Glp1r* expression in metabolic and non-metabolic tissues appears to be crucial to maintain glucose homeostasis. In addition, the identification of the signaling pathways that regulate *Glp1r* expression could represent a new target towards T2D treatment. For instance, Pax6 (64) or farnesoid X receptor (65) directly modulate the *Glp1r* gene expression at chromatin level, thus targeting these pathways might be promising to improve glucose homeostasis and insulin secretion in a metabolic stress context. We also observe that overexpressing *E2F1* in pancreatic β cells increases *Glp1r* expression and improves glucose tolerance upon high fat diet, suggesting that targeting this pathway would also represent an alternative strategy, with caution regarding the proliferative functions displayed by the E2F1/pRb pathway. Moreover, since *E2F1* expression is decreased in T2D islets (37), our study suggests a putative association between the loss of *GLP1R* expression in T2D patients and the loss of *E2F1* expression. Although single cell RNA sequencing analysis are required to unequivocally demonstrate concomitant decreased levels of *E2F1* and *GLP1R* within the same cell, we can speculate that the long term loss of efficacy of Glp1r agonist treatment to restore glucose homeostasis in T2D patients may be related, at least partially, to impaired E2F1-dependent modulation of *GLP1R* expression.

Following pioneer observations that the E2F1-CDK4-pRb pathway is regulated by glucose and insulin in pancreatic islets (17), we show here that the treatment of pancreatic islets with a Glp1r agonist modulates E2F1 pathway through the phosphorylation of pRb and the activation of the E2F1 transcriptional activity. Our data are in line with previous studies showing that the treatment of Ins1 cells with exendin-4 decreases both mRNA and protein levels of pRb and increases E2f1 protein levels (53). It is well established that exendin-4 and Glp1r agonists increases cell cycle progression through the increased expression of cyclins A2 and cyclin D1, stimulating pancreatic β-cell proliferation (66, 67). Since the E2F1-pRb-CDK4 pathway positively modulates the expression of *Ccna2* and *Ccnd1* (68), our data suggest that the cell cycle progression of pancreatic β cell through the induction of *Ccna2* and *Ccnd1* with exendin-4 treatment could be associated with an increase of E2f1 expression and/or activity. Moreover, in human islets, the treatment with exendin-4 also increases the expression of several cyclins, leading to increased pancreatic β-cell proliferation (69). Here we observed an increase of pRb phosphorylation in pancreatic islets treated with exendin-4 and a raise in *E2f1* expression in Min6 cells treated with exendin-4, suggesting that β-cell proliferation induced by Glp1r agonist could be under the control by the E2F1-pRb pathway.

A limitation of this study is the use of the RIP-Cre mice (39) which allows Cre-recombinase expression at the early stage of pancreatic β-cell differentiation, between E15.5 and E18.5 (70). Since E2f1 plays a key role in both post-natal proliferation of β cells (18) and early pancreas development by targeting *Pdx1* and *Ngn3* expression through the Cdk4 pathway (19), the deletion of the *E2f1* gene using Rip-Cre mice may affect early β-cell differentiation and mass. Therefore, we cannot exclude that the early loss of *E2f1* expression could contribute to decreased *Glp1r* expression in *E2f1-* deficient pancreatic islets during pancreatic β-cell development. The use of an inducible model, such as the Mip-Cre^ERT^ mice (71), may allow to determine whether the deletion of the *E2f1* gene in non-proliferating adult β cells could also result in impaired glucose homeostasis related to the modulation of *Glp1r* expression by E2F1 in adulthood.

In summary, our study reveals a crosstalk between E2f1 and Glp-1 signaling pathways in both mice and human pancreatic islets. It opens the possibility to understanding the differential effects of Glp-1 in the control of β-cell mass and function and their relevant molecular mechanisms. How this crosstalk modulates β-cell proliferation, which downstream signaling molecules are involved, and how these mechanisms are actionable in T2D pathophysiology remain open questions that require further investigations.

## Author contributions

C.B., E.Co., G.P., X.G., N.R., C.C., M.M., R.B., L.R. and E.Ca. contributed to the *in vivo* and cellular experiments. Z.B., P.M., J.K.C., F.P., P.F., A.B. and F.O. provided reagents (J.K.C. and F.P. provided human islets) and data and discussed the results from the study. J.-S.A. designed the study, supervised the project and contributed to experiments and/or their analysis and the funding of this project. C.B., E.C. and J.-S.A. wrote and/or edited the manuscript. J.-S.A. is the guarantor of this work and, as such, had full access to all of the data in the study and take responsibility for the integrity of the data and the accuracy of the data analysis.

## Acknowledgements

We thank Dr David Blum, Dr Stéphane Dalle and Dr Benoit Pourcet and members of the INSERM U1283/CNRS UMR 8199/EGID for helpful discussions and critical reading of our manuscript. Human islets were provided through the JDRF award 31-2008-416 (ECIT Islet for Basic Research program). The authors thank UMS2014-US41 and the Experimental Resources platform from Université de Lille, especially Cyrille Degraeve, Mélanie Besegher and Julien Devassine for animal care. We thank the Department of Histology from the Lille Medicine Faculty, particularly H. Gevaert and R.M. Siminski, for histological preparations. This work was supported by the Agence Nationale de la Recherche (ANR) grants « European Genomic Institute for Diabetes » E.G.I.D, ANR-10-LABX-46 and Equipex 2010 ANR-10-EQPX-07-01; ‘LIGAN-PM’ Genomics platform, a French State fund managed by ANR under the frame program Investissements d’Avenir I-SITE ULNE / ANR-16-IDEX-0004 ULNE (to P.F., A.B. and J-S.A), ANR BETAPLASTICITY (ANR-17-CE14-0034 to J-S.A.), European Foundation for the Study of Diabetes (EFSD, to J-S.A.), INSERM, CNRS, Institut Pasteur de Lille (CPER CTRL Melodie, to E.C. and J-S.A.), Association pour la Recherche sur le Diabète (to J-S.A.), Université de Lille (to C.B., F.O., X.G. and J-S.A.), I-SITE ULNE (EpiRNAdiab Sustain grant to J-S.A.), Conseil Régional Hauts de France and Métropole Européenne de Lille (to J-S.A.), F.E.D.E.R. (Fonds Européen de Développement Régional, to P.F. and J-S.A.) and Société Francophone du Diabète (to J-S.A).

## References

1. Weir GC, and Bonner-Weir S. Five stages of evolving beta-cell dysfunction during progression to diabetes. Diabetes. 2004;53 Suppl 3:S16–21.

2. Drucker DJ. Mechanisms of Action and Therapeutic Application of Glucagon-like Peptide-1. Cell Metab. 2018;27(4):740–56.

3. Holst JJ, Orskov C, Nielsen OV, and Schwartz TW. Truncated glucagon-like peptide I, an insulin-releasing hormone from the distal gut. FEBS Lett. 1987;211(2):169–74.

4. Chambers AP, Sorrell JE, Haller A, Roelofs K, Hutch CR, Kim KS, et al. The Role of Pancreatic Preproglucagon in Glucose Homeostasis in Mice. Cell Metab. 2017;25(4):927–34 e3.

5. Thorens B. Expression cloning of the pancreatic beta cell receptor for the gluco-incretin hormone glucagon-like peptide 1. Proc Natl Acad Sci U S A. 1992;89(18):8641–5.

6. Nauck MA, Quast DR, Wefers J, and Meier JJ. GLP-1 receptor agonists in the treatment of type 2 diabetes - state-of-the-art. Mol Metab. 2020:101102.

7. Baggio LL, and Drucker DJ. Biology of incretins: GLP-1 and GIP. Gastroenterology. 2007;132(6):2131–57.

8. Holz GGt, Kuhtreiber WM, and Habener JF. Pancreatic beta-cells are rendered glucose-competent by the insulinotropic hormone glucagon-like peptide-1(7-37). Nature. 1993;361(6410):362–5.

9. Kreymann B, Williams G, Ghatei MA, and Bloom SR. Glucagon-like peptide-1 7-36: a physiological incretin in man. Lancet. 1987;2(8571):1300–4.

10. de Heer J, Rasmussen C, Coy DH, and Holst JJ. Glucagon-like peptide-1, but not glucose-dependent insulinotropic peptide, inhibits glucagon secretion via somatostatin (receptor subtype 2) in the perfused rat pancreas. Diabetologia. 2008;51(12):2263–70.

11. Cornu M, Modi H, Kawamori D, Kulkarni RN, Joffraud M, and Thorens B. Glucagon-like peptide-1 increases beta-cell glucose competence and proliferation by translational induction of insulin-like growth factor-1 receptor expression. J Biol Chem. 2010;285(14):10538–45.

12. Friedrichsen BN, Neubauer N, Lee YC, Gram VK, Blume N, Petersen JS, et al. Stimulation of pancreatic beta-cell replication by incretins involves transcriptional induction of cyclin D1 via multiple signalling pathways. J Endocrinol. 2006;188(3):481–92.

13. Liu Z, and Habener JF. Glucagon-like peptide-1 activation of TCF7L2-dependent Wnt signaling enhances pancreatic beta cell proliferation. J Biol Chem. 2008;283(13):8723–35.

14. Quoyer J, Longuet C, Broca C, Linck N, Costes S, Varin E, et al. GLP-1 mediates antiapoptotic effect by phosphorylating Bad through a beta-arrestin 1-mediated ERK1/2 activation in pancreatic beta-cells. J Biol Chem. 2010;285(3):1989–2002.

15. Li Y, Hansotia T, Yusta B, Ris F, Halban PA, and Drucker DJ. Glucagon-like peptide-1 receptor signaling modulates beta cell apoptosis. J Biol Chem. 2003;278(1):471–8.

16. Buteau J, El-Assaad W, Rhodes CJ, Rosenberg L, Joly E, and Prentki M. Glucagon-like peptide-1 prevents beta cell glucolipotoxicity. Diabetologia. 2004;47(5):806–15.

17. Annicotte JS, Blanchet E, Chavey C, Iankova I, Costes S, Assou S, et al. The CDK4-pRB-E2F1 pathway controls insulin secretion. Nat Cell Biol. 2009;11(8):1017–23.

18. Fajas L, Annicotte JS, Miard S, Sarruf D, Watanabe M, and Auwerx J. Impaired pancreatic growth, beta cell mass, and beta cell function in E2F1 (−/−)mice. J Clin Invest. 2004;113(9):1288–95.

19. Kim SY, and Rane SG. The Cdk4-E2f1 pathway regulates early pancreas development by targeting Pdx1+ progenitors and Ngn3+ endocrine precursors. Development. 2011;138(10):1903–12.

20. Dimova DK, and Dyson NJ. The E2F transcriptional network: old acquaintances with new faces. Oncogene. 2005;24(17):2810–26.

21. Mendoza PR, and Grossniklaus HE. The Biology of Retinoblastoma. Prog Mol Biol Transl Sci. 2015;134:503–16.

22. Flemington EK, Speck SH, and Kaelin WG, Jr. E2F-1-mediated transactivation is inhibited by complex formation with the retinoblastoma susceptibility gene product. Proc Natl Acad Sci U S A. 1993;90(15):6914–8.

23. Blais A, and Dynlacht BD. E2F-associated chromatin modifiers and cell cycle control. Curr Opin Cell Biol. 2007;19(6):658–62.

24. Kato J, Matsushime H, Hiebert SW, Ewen ME, and Sherr CJ. Direct binding of cyclin D to the retinoblastoma gene product (pRb) and pRb phosphorylation by the cyclin D-dependent kinase CDK4. Genes Dev. 1993;7(3):331–42.

25. Schulze A, Zerfass K, Spitkovsky D, Henglein B, and Jansen-Durr P. Activation of the E2F transcription factor by cyclin D1 is blocked by p16INK4, the product of the putative tumor suppressor gene MTS1. Oncogene. 1994;9(12):3475–82.

26. Poppy Roworth A, Ghari F, and La Thangue NB. To live or let die - complexity within the E2F1 pathway. Mol Cell Oncol. 2015;2(1):e970480.

27. Denechaud PD, Fajas L, and Giralt A. E2F1, a Novel Regulator of Metabolism. Frontiers in endocrinology. 2017;8:311.

28. Kahoul Y, Oger F, Montaigne J, Froguel P, Breton C, and Annicotte JS. Emerging Roles for the INK4a/ARF (CDKN2A) Locus in Adipose Tissue: Implications for Obesity and Type 2 Diabetes. Biomolecules. 2020;10(9).

29. Fajas L, Landsberg RL, Huss-Garcia Y, Sardet C, Lees JA, and Auwerx J. E2Fs regulate adipocyte differentiation. Dev Cell. 2002;3(1):39–49.

30. Chen J, Yang Y, Li S, Yang Y, Dai Z, Wang F, et al. E2F1 Regulates Adipocyte Differentiation and Adipogenesis by Activating ICAT. Cells. 2020;9(4).

31. Denechaud PD, Lopez-Mejia IC, Giralt A, Lai Q, Blanchet E, Delacuisine B, et al. E2F1 mediates sustained lipogenesis and contributes to hepatic steatosis. J Clin Invest. 2016;126(1):137–50.

32. Giralt A, Denechaud PD, Lopez-Mejia IC, Delacuisine B, Blanchet E, Bonner C, et al. E2F1 promotes hepatic gluconeogenesis and contributes to hyperglycemia during diabetes. Mol Metab. 2018;11:104–12.

33. Lai Q, Giralt A, Le May C, Zhang L, Cariou B, Denechaud PD, et al. E2F1 inhibits circulating cholesterol clearance by regulating Pcsk9 expression in the liver. JCI Insight. 2017;2(10).

34. Blanchet E, Annicotte JS, Lagarrigue S, Aguilar V, Clape C, Chavey C, et al. E2F transcription factor-1 regulates oxidative metabolism. Nat Cell Biol. 2011;13(9):1146–52.

35. Blanchet E, Annicotte JS, Pradelli LA, Hugon G, Matecki S, Mornet D, et al. E2F transcription factor-1 deficiency reduces pathophysiology in the mouse model of Duchenne muscular dystrophy through increased muscle oxidative metabolism. Hum Mol Genet. 2012;21(17):3910–7.

36. Grouwels G, Cai Y, Hoebeke I, Leuckx G, Heremans Y, Ziebold U, et al. Ectopic expression of E2F1 stimulates beta-cell proliferation and function. Diabetes. 2010;59(6):1435–44.

37. Lupi R, Mancarella R, Del Guerra S, Bugliani M, Del Prato S, Boggi U, et al. Effects of exendin-4 on islets from type 2 diabetes patients. Diabetes Obes Metab. 2008;10(6):515–9.

38. Rabhi N, Denechaud PD, Gromada X, Hannou SA, Zhang H, Rashid T, et al. KAT2B Is Required for Pancreatic Beta Cell Adaptation to Metabolic Stress by Controlling the Unfolded Protein Response. Cell Rep. 2016;15(5):1051–61.

39. Herrera PL, Orci L, and Vassalli JD. Two transgenic approaches to define the cell lineages in endocrine pancreas development. Mol Cell Endocrinol. 1998;140(1-2):45–50.

40. Scheijen B, Bronk M, and Meer TVD. Constitutive E2F1 Overexpression Delays Endochondral Bone Formation by Inhibiting Chondrocyte Differentiation. Society. 2003;23(10):3656–68.

41. Kerr-Conte J, Vandewalle B, Moerman E, Lukowiak B, Gmyr V, Arnalsteen L, et al. Upgrading pretransplant human islet culture technology requires human serum combined with media renewal. Transplantation. 2010;89(9):1154–60.

42. Rosales-Hurtado M, Lebeau A, Bourouh C, Cebrian-Torrejon G, Albalat M, Jean M, et al. Improved synthesis, resolution, absolute configuration determination and biological evaluation of HLM006474 enantiomers. Bioorg Med Chem Lett. 2019;29(3):380–2.

43. Rabhi N, Hannou SA, Gromada X, Salas E, Yao X, Oger F, et al. Cdkn2a deficiency promotes adipose tissue browning. Mol Metab. 2018;8:65–76.

44. Lee C, and Huang CH. LASAGNA-Search: an integrated web tool for transcription factor binding site search and visualization. Biotechniques. 2013;54(3):141–53.

45. Rabhi N, Denechaud PD, Gromada X, Hannou SA, Zhang H, Rashid T, et al. KAT2B is required for pancreatic beta cell adaptation to metabolic stress by controlling the unfolded protein response signaling. Cell Reports. 2016;In press.

46. Muller TD, Finan B, Bloom SR, D’Alessio D, Drucker DJ, Flatt PR, et al. Glucagon-like peptide 1 (GLP-1). Mol Metab. 2019;30:72–130.

47. Ashcroft SJ. Protein phosphorylation and beta-cell function. Diabetologia. 1994;37 Suppl 2:S21–9.

48. Jhala US, Canettieri G, Screaton RA, Kulkarni RN, Krajewski S, Reed J, et al. cAMP promotes pancreatic beta-cell survival via CREB-mediated induction of IRS2. Genes Dev. 2003;17(13):1575–80.

49. Fujitani Y. Transcriptional regulation of pancreas development and beta-cell function [Review]. Endocr J. 2017;64(5):477–86.

50. Wang X, Zhou J, Doyle ME, and Egan JM. Glucagon-like peptide-1 causes pancreatic duodenal homeobox-1 protein translocation from the cytoplasm to the nucleus of pancreatic beta-cells by a cyclic adenosine monophosphate/protein kinase A-dependent mechanism. Endocrinology. 2001;142(5):1820–7.

51. Kodama S, Toyonaga T, Kondo T, Matsumoto K, Tsuruzoe K, Kawashima J, et al. Enhanced expression of PDX-1 and Ngn3 by exendin-4 during beta cellregeneration in STZ-treated mice. Biochem Biophys Res Commun. 2005;327(4):1170–8.

52. Cam H, and Dynlacht BD. Emerging roles for E2F: beyond the G1/S transition and DNA replication. Cancer Cell. 2003;3(4):311–6.

53. Cai EP, Luk CT, Wu X, Schroer SA, Shi SY, Sivasubramaniyam T, et al. Rb and p107 are required for alpha cell survival, beta cell cycle control and glucagon-like peptide-1 action. Diabetologia. 2014;57(12):2555–65.

54. Cai EP, Wu X, Schroer SA, Elia AJ, Nostro MC, Zacksenhaus E, et al. Retinoblastoma tumor suppressor protein in pancreatic progenitors controls alpha-and beta-cell fate. Proc Natl Acad Sci U S A. 2013;110(36):14723–8.

55. Boni-Schnetzler M, Hauselmann SP, Dalmas E, Meier DT, Thienel C, Traub S, et al. beta Cell-Specific Deletion of the IL-1 Receptor Antagonist Impairs beta Cell Proliferation and Insulin Secretion. Cell Rep. 2018;22(7):1774–86.

56. Hinnen D. Glucagon-Like Peptide 1 Receptor Agonists for Type 2 Diabetes. Diabetes Spectr. 2017;30(3):202–10.

57. Campbell JE, and Drucker DJ. Pharmacology, physiology, and mechanisms of incretin hormone action. Cell Metab. 2013;17(6):819–37.

58. Ahren B. Incretin dysfunction in type 2 diabetes: clinical impact and future perspectives. Diabetes Metab. 2013;39(3):195–201.

59. Knop FK, Vilsboll T, Hojberg PV, Larsen S, Madsbad S, Volund A, et al. Reduced incretin effect in type 2 diabetes: cause or consequence of the diabetic state? Diabetes. 2007;56(8):1951–9.

60. Pan QR, Li WH, Wang H, Sun Q, Xiao XH, Brock B, et al. Glucose, metformin, and AICAR regulate the expression of G protein-coupled receptor members in INS-1 beta cell. Horm Metab Res. 2009;41(11):799–804.

61. Xu G, Kaneto H, Laybutt DR, Duvivier-Kali VF, Trivedi N, Suzuma K, et al. Downregulation of GLP-1 and GIP receptor expression by hyperglycemia: possible contribution to impaired incretin effects in diabetes. Diabetes. 2007;56(6):1551–8.

62. Kimura T, Obata A, Shimoda M, Shimizu I, da Silva Xavier G, Okauchi S, et al. Down-regulation of vascular GLP-1 receptor expression in human subjects with obesity. Sci Rep. 2018;8(1):10644.

63. Rajan S, Dickson LM, Mathew E, Orr CM, Ellenbroek JH, Philipson LH, et al. Chronic hyperglycemia downregulates GLP-1 receptor signaling in pancreatic beta-cells via protein kinase A. Mol Metab. 2015;4(4):265–76.

64. Gosmain Y, Katz LS, Masson MH, Cheyssac C, Poisson C, and Philippe J. Pax6 is crucial for beta-cell function, insulin biosynthesis, and glucose-induced insulin secretion. Mol Endocrinol. 2012;26(4):696–709.

65. Kong X, Feng L, Yan D, Li B, Yang Y, and Ma X. FXR-mediated epigenetic regulation of GLP-1R expression contributes to enhanced incretin effect in diabetes after RYGB. J Cell Mol Med. 2021.

66. Kim MJ, Kang JH, Park YG, Ryu GR, Ko SH, Jeong IK, et al. Exendin-4 induction of cyclin D1 expression in INS-1 beta-cells: involvement of cAMP-responsive element. J Endocrinol. 2006;188(3):623–33.

67. Song WJ, Schreiber WE, Zhong E, Liu FF, Kornfeld BD, Wondisford FE, et al. Exendin-4 stimulation of cyclin A2 in beta-cell proliferation. Diabetes. 2008;57(9):2371–81.

68. Stanelle J, Stiewe T, Theseling CC, Peter M, and Putzer BM. Gene expression changes in response to E2F1 activation. Nucleic Acids Res. 2002;30(8):1859–67.

69. Dai C, Hang Y, Shostak A, Poffenberger G, Hart N, Prasad N, et al. Age-dependent human beta cell proliferation induced by glucagon-like peptide 1 and calcineurin signaling. J Clin Invest. 2017;127(10):3835–44.

70. Herrera PL, Huarte J, Sanvito F, Meda P, Orci L, and Vassalli JD. Embryogenesis of the murine endocrine pancreas; early expression of pancreatic polypeptide gene. Development. 1991;113(4):1257–65.

71. Tamarina NA, Roe MW, and Philipson L. Characterization of mice expressing Ins1 gene promoter driven CreERT recombinase for conditional gene deletion in pancreatic beta-cells. Islets. 2014;6(1):e27685.

