## Supplementary information for "A crosstalk between E2F1 and GLP-1 signaling pathways modulates insulin secretion"

**Bourouh *et al.*, Supplentary Figure S1**

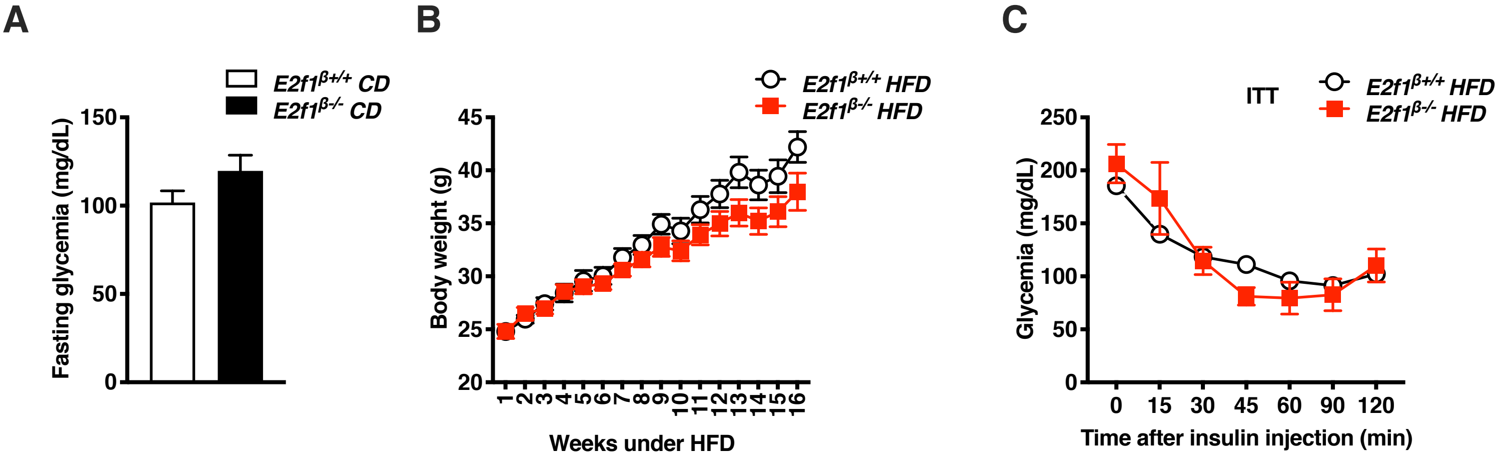

Bourouh *et al.*, Supplentary Figure S2

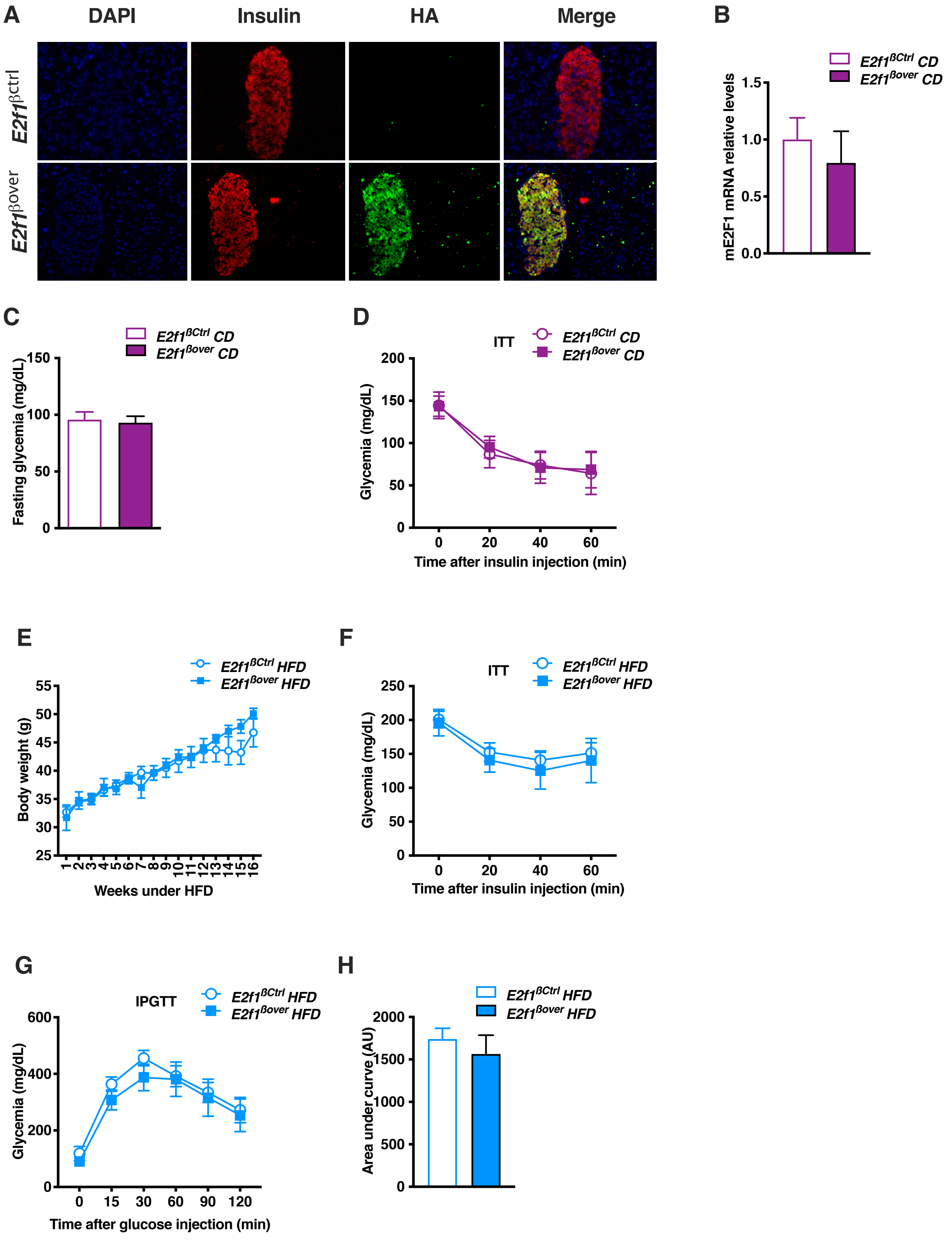

**A**

### E2F1 binding motif

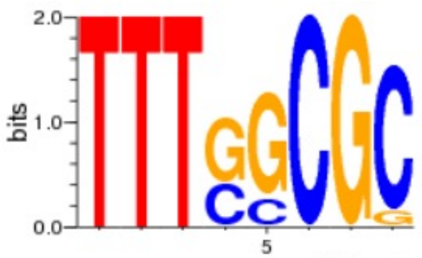


**B**

### E2f binding sites in the mouse *Glp1r* promoter

| Sequence | Position (0-based) | Strand | P-value |
|---|---|---|---|
| TTCGGCGC | 797 | - | 0.002275 |
| TTCGGCG | 798 | - | 0.0299 |
| TCTGGCCC | 835 | + | 0.037075 |
| TTGGGTGC | 483 | - | 0.037075 |
| TGTGGCTC | 34 | - | 0.037075 |
| TCTGGAGC | 162 | + | 0.037075 |
| TGTGGTGC | 723 | - | 0.037075 |

A

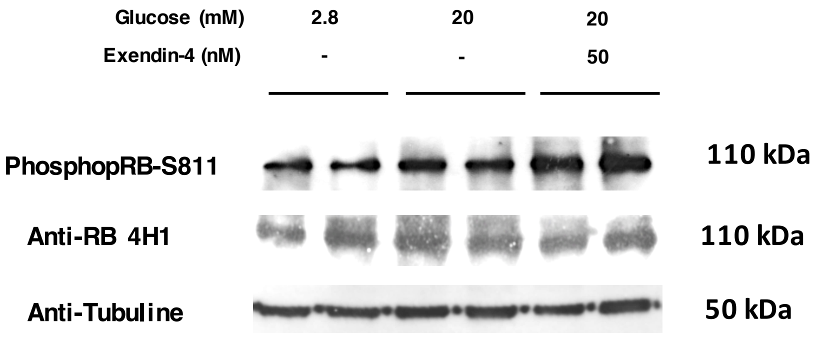
Glucose (mM)
2.8
20
20
Exendin-4 (nM)
-
-
50
PhosphopRB-S811
110 kDa
Anti-RB 4H1
110 kDa
Anti-Tubuline
50 kDa

B

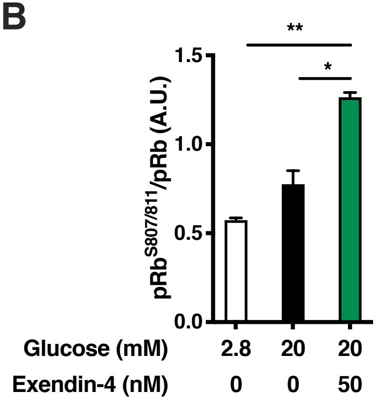
pRb$^{S807/811}$/pRb (A.U.)
1.5
1.0
0.5
0.0
**
*
Glucose (mM)
2.8
20
20
Exendin-4 (nM)
0
0
50

C

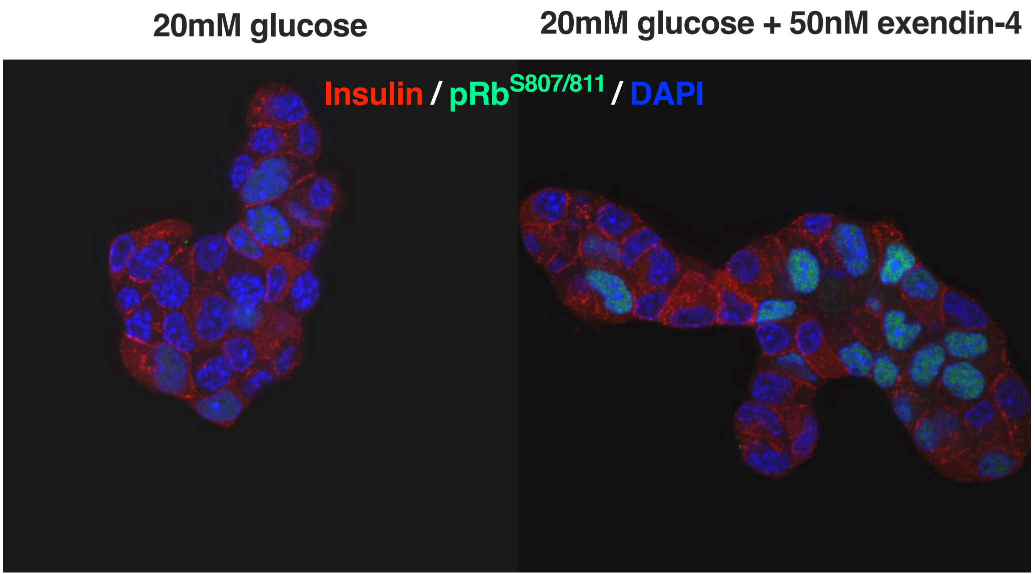
20mM glucose
20mM glucose + 50nM exendin-4
Insulin / pRb$^{S807/811}$ / DAPI

**Supplementary information**

**Supplementary Figure S1, related to Figure 1. Metabolic parameters of *E2f1*$^{\beta+/+}$ and *E2f1*β-/- mice. (A)** Blood glucose level of *E2f1*$^{\beta+/+}$ and *E2f1*$^{\beta-/-}$ mice fed with a CD after 16h of fasting **(B)** Evolution of body weight of *E2f1*$^{\beta+/+}$ and *E2f1*$^{\beta-/-}$ mice fed with a high fat diet (n=7). **(C)** Blood glucose levels during intraperitoneal insulin tolerance test (ITT) from *E2f1*$^{\beta+/+}$ and *E2f1*$^{\beta-/-}$ mice after 16 weeks of HFD (*n*=5). All values are expressed as mean ± s.e.m. and were analyzed by two-tailed unpaired *t*-test (A) or two-way ANOVA followed by a Tukey's *post-hoc* test (B, C).

**Supplementary Figure S2, related to Figure 2. Characterization of *E2f1*$^{\beta ctrl}$ *E2f1*$^{\beta over}$ mice. (A)** Immunofluorescence analysis of pancreatic sections showing a staining for insulin (red), HA (green) and DAPI nuclear staining (blue) on representative sections from pancreas of *E2f1*$^{\beta ctrl}$ and *E2f1*$^{\beta over}$. **(B)** *mE2f1* mRNA expression levels from *E2f1*$^{\beta ctrl}$ and *E2f1*$^{\beta over}$ mice (*n=6).* **(C)** Fasting blood glucose levels of *E2f1*$^{\beta ctrl}$ and *E2f1*$^{\beta over}$ mice (n=10-14). (**D)** Blood glucose levels during intraperitoneal insulin tolerance test (ITT) from *E2f1*$^{\beta ctrl}$ and *E2f1*$^{\beta over}$ mice fed with CD (*n*=5). **(E)** Body weight of *E2f1*$^{\beta ctrl}$ and *E2f1*$^{\beta over}$ mice fed with a high fat diet for 16 weeks (n=5). **(F)** ITT was carried out on *E2f1*$^{\beta ctrl}$ and *E2f1*$^{\beta over}$ mice fed with HFD for 17 weeks (*n*=5). **(G)** IPGTT was performed after 16h fasting in *E2f1*$^{\beta ctrl}$ and *E2f1*$^{\beta over}$ mice fed with HFD for 12 weeks (*n*=5). (**H**) Area under curve of IPGTT from *E2f1*$^{\beta ctrl}$ and *E2f1*$^{\beta over}$ mice. All values are expressed as mean ± s.e.m. and were analyzed by two-tailed unpaired *t*-test (B, C, H) or two-way ANOVA followed by a Tukey's *post-hoc* test (D, E, F, G).

**Supplementary Figure S3, related to Figure 3. E2f1 binding sites motif in the mouse *Glp1r* promoter. (A)** The consensus motif of E2F1 DNA binding site is represented. **(B)** *In silico* analysis of the mouse *Glp1r* promoter using LASAGNA identifying E2f1 binding sites present in the promoter region of the mouse *Glp1r* promoter.

**Supplementary Figure S4, related to figure 4. Exendin-4 treatment induces pRb phosphorylation at serine 807/811. (A)** Western blot assay of total pRb and phosphorylated pRb$^{S807/811}$ in Min6 exposed to 2.8 mM glucose, 20 mM glucose or to

50 nM exendin-4 during 30 minutes. Tubulin was used as loading control. **(B)** Quantification of pRb$^{S807/811}$ protein levels. The western blot band instensty of phosphorylated and total pRb was calculated using ImageJ software and the ratio between phosphorylated pRB (pRb$^{S807/811}$) and total pRB is represented. (*n*=3) **(C)** Immunofluorescence analysis of phosphorylated pRb$^{S807/811}$ (green), insulin (red) and DAPI nuclear staining (blue) in Min6 cells exposed to 20 mM glucose and treated or not with 50 nM of exendin-4 during 30 minutes.

**Supplementary Table S1: List of oligonucleotides**

| Gene name | Gene Symbol | Species | Primer |
|---|---|---|---|
| *Cyclophillin* | *Cyclo* | *Mouse* | *ATGGCACTGGCGGCAGGTCC* |
| | | | *TTGCCATTCCTGGACCCAAA* |
| *E2f transcription factor 1* | *E2f1* | *Mouse* | *GTGAATCGATAGTACTAACATACG* |
| | | | *GTATCTCTTCATAGCCTTATGCAG* |
| *E2f transcription factor 1 Exon3* | *E2f1* | *Mouse* | ACAGCTGCAACTGCTTTCGGAG |
| | | | AGCTTGTAGTTGGGTCTCAGGAGG |
| *Glucagon-like peptide 1 receptor* | *Glp1r* | *Mouse* | *GTTTCCTCACGGAAGCGCCA* |
| | | | *AAGGAACCTGGGGGCCCATC* |
| *Pancreatic and duodenal homeobox1* | *Pdx1* | *Mouse* | *ATTGTGCGGTGACCTCGGGC* |
| | | | *GATGCTGGAGGGCTGTGGCG* |
| *Glucagon-like peptide 1 receptor promotor* | *Glp1r-prom* | *Mouse* | *CTGCGGCTCTTAAACCTGAG* |
| | | | *CTGTCCTCTTCCGCCTGG* |
| *Cyclin A2* | *Ccna2* | *Mouse* | *GGCTGACACTCTTTCCG* |
| | | | *CTGGTAGCAAGAATTAGAGCAT* |
| *Cyclophillin* | *hCyclo* | *Human* | *ATGGCACTGGTGGCAAGTCC* |
| | | | *TTGCCATTCCTGGACCCAAA* |
| *E2f transcription factor 1* | *hE2f1* | *Human* | *GAGAAGTCCTCCCGCACATG* |
| | | | *CACAGATCCCAGCCAGTCTC* |
| *Glucagon-like peptide 1 receptor* | *hGlp1r* | *Human* | *AGTCCAAGCGAGGGGAAAGA* |
| | | | *GAGGCGATAACCAGAGCAGAG* |

### Supplementary Table S2: Donor information.

| **Islet preparation** | **1** | **2** | **3** |
|---|---|---|---|
| **MANDATORY INFORMATION** | | | |
| Unique identifier | H1032 | H1039 | H1041 |
| Donor age (years) | 35 | 54 | 37 |
| Donor sex (M/F) | M | F | F |
| Donor BMI (kg/m$^2$) | 35.2 | 24.2 | 24.3 |
| Donor $HbA_{1c}$ or other measure of blood glucose control | 5.4 | 6.3 | 5.8 |
| Origin/source of islets[b] | ECIT | ECIT | ECIT |
| Islet isolation centre | LILLE | LILLE | LILLE |
| Donor history of diabetes? Please select yes/no from drop down list | No | No | No |
| **If Yes, complete the next two lines if this information is available** | | | |
| Diabetes duration (years) | NA | NA | NA |
| Glucose-lowering therapy at time of death[c] | NA | NA | NA |
| **RECOMMENDED INFORMATION** | | | |
| Donor cause of death | Stroke | Stroke | Anoxia |
| Warm ischaemia time (h) | No | No | No |
| Cold ischaemia time (h) | 4h34 | 6h22 | 5h30 |
| Estimated purity (%) | 80 | 80 | 70 |
| Estimated viability (%) | 95.5 | 94.7 | 93.3 |
| Total culture time (h)[d] | 18 | 18 | 19 |
| Glucose-stimulated insulin secretion or other functional measurement[e] | 7,19 | 3,04 | N/A |
| Handpicked to purity? Please select yes/no from drop down list | No | No | No |
